# The transcriptional legacy of developmental stochasticity

**DOI:** 10.1101/2019.12.11.873265

**Authors:** Sara Ballouz, Maria T. Pena, Frank M. Knight, Linda B. Adams, Jesse A. Gillis

## Abstract

Genetic variation, epigenetic regulation and major environmental stimuli are key contributors to phenotypic variation, but the influence of minor perturbations or “noise” has been difficult to assess in mammals. In this work, we uncover one major axis of random variation with a large and permanent influence: developmental stochasticity. By assaying the transcriptome of wild monozygotic quadruplets of the nine-banded armadillo, we find that persistent changes occur early in development, and these give rise to clear transcriptional signatures which uniquely characterize individuals relative to siblings. Comparing these results to human twins, we find the transcriptional signatures which define individuals exhibit conserved co-expression, suggesting a substantial fraction of phenotypic and disease discordance within mammals arises from developmental stochasticity.

**One sentence summary:** Longitudinal gene expression in identical armadillo quadruplets reveals a major role for developmental stochasticity.

## Main text

Variability in human phenotype is the product of genetic and environmental contributions, along with a complex interplay between the two (*1*). While genomic data has permitted valuable progress in our understanding of both heritable and non-heritable phenotypic variation, this progress has been more piecemeal in sources of non-heritable variation. All studies of genetic or environmental influences on phenotype are affected by this unexplained, non-heritable variability or ‘noise’ (*2*). One possibility is that ‘noise’ can be partitioned into well-defined categories of its own, based on underlying mechanisms. Development has long been thought to be a potential driver of unexplained phenotypic variability (*3*); it is a time when small initial changes can permanently propagate forward to large later effect (**Fig. 1A**). While programmatic variability in development has received particular attention (*4, 5*), early random effects could be a major source of phenotypic variance (**Fig. 1B**). In order to measure this developmental stochasticity, tight environmental and genetic are necessary to minimize external drivers of variability, while outbred genetics are necessary to maximize the likely functional implications of observed variability. In this work, we exploit polyembrony in *Dasypus novemcinctus* (the nine-banded armadillo) to control for genetics and environment and quantify the impact of early random variation across outbred genetic backgrounds (**Fig. 1C-D**). Uniquely among mammals, armadillos have evolved a reproductive strategy that produces litters of identical quadruplets, creating a unique opportunity for outbred control of both genetics and environment.

**Fig. 1:**
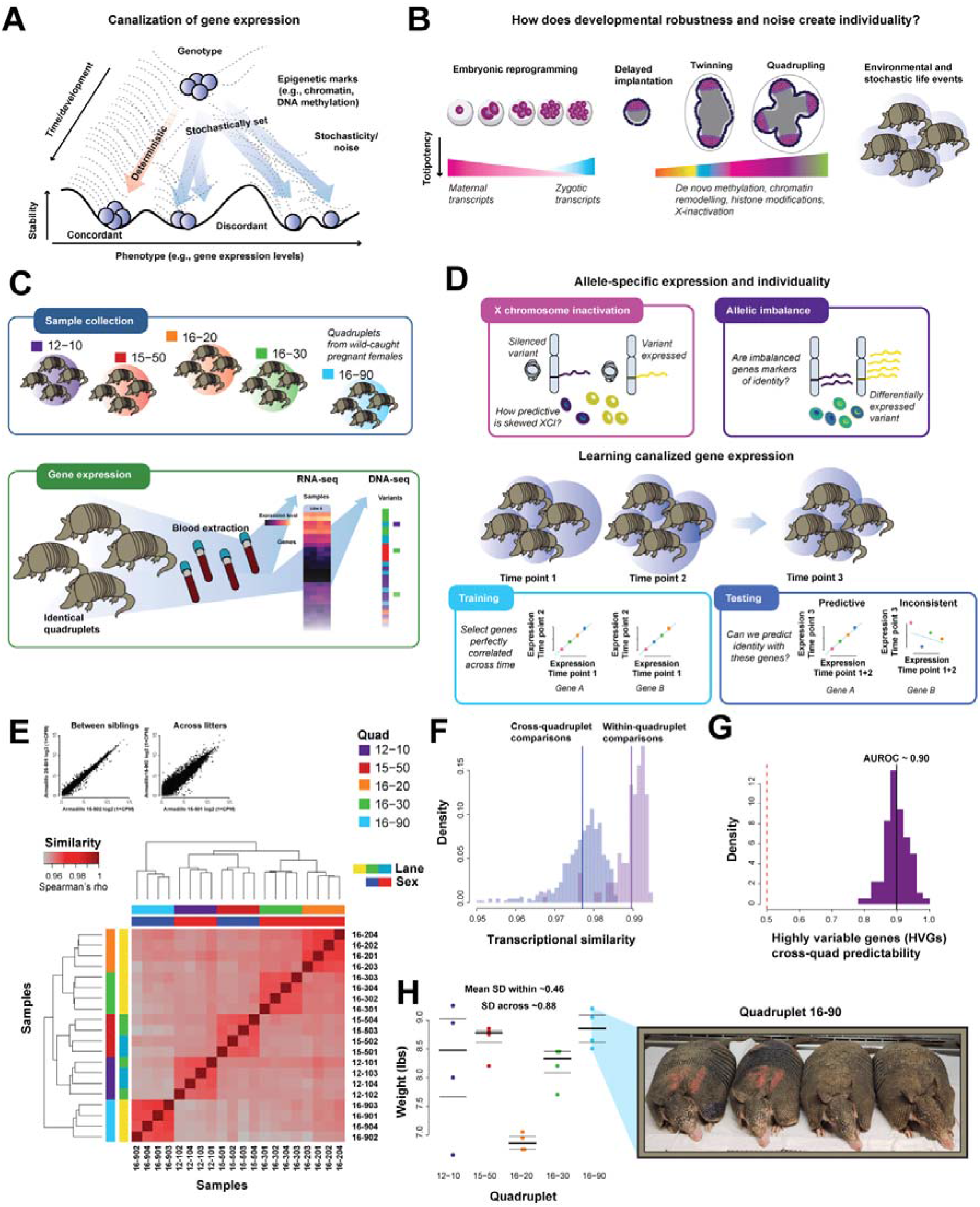
Defining transcriptional identity. (A) Gene expression is canalized. (B) This can occur at different stages of development. (C) The study design. (D) Measuring identity. (E) Transcriptional similarity is higher between siblings than across armadillo quadruplets. Heatmap of sample-sample correlations for the first time point with example sample-sample scatterplots. (F) Distributions of the within and between correlations. (G) Variable genes within a quad are variable in other quadruplets. (H) Weight similarity showing lower variance within than across all, and a set of quadruplets.

Our central experimental strategy is to measure gene expression over time and look for signatures permanently distinguishing siblings from one another. As an aggregate readout of epigenetic variability between the genetically identical individuals, gene expression serves as a likely intermediate to capture variability with potential phenotypic impact. Because the environment is controlled, armadillo expression profiles are extremely similar even across genetic background (**Fig. 1E**, r>0.95). Within sibling cohorts, the correlations are tighter (r>0.98) reflecting the expected addition of controlled genetics (**Fig. 1F**). However, the genes which do vary in their expression across individuals within each quad are consistent and can be predicted from the variability in the other quads (**Fig. 1G**, AUROC~0.9).

Allelic expression imbalance on the X-chromosome in females is particularly striking in its variability. As X-chromosome inactivation (XCI) is stochastic when it first occurs (*6*) and then is maintained in cell lineages (*7*), it creates permanent variability between individuals (e.g., **Fig. 2C**). The choice of which X to inactivate is effectively a coin flip at the time of inactivation, meaning that the population variance is defined from a simple binomial model (*8*). Because this variability is permanently fixed within cell lineages, it does not change later in development and the observed variance provides an estimate of the number of cells present in the population at the time of X-inactivation. In our female samples, we can estimate XCI occurring after the embryos have split, at a population size of 25 cells (**Fig. 2D**), +/− ~1 cell division. This occurs after the developing armadillo embryo has split into four, causing each sibling to be a distinct individual with respect to XCI.

**Fig. 2:**
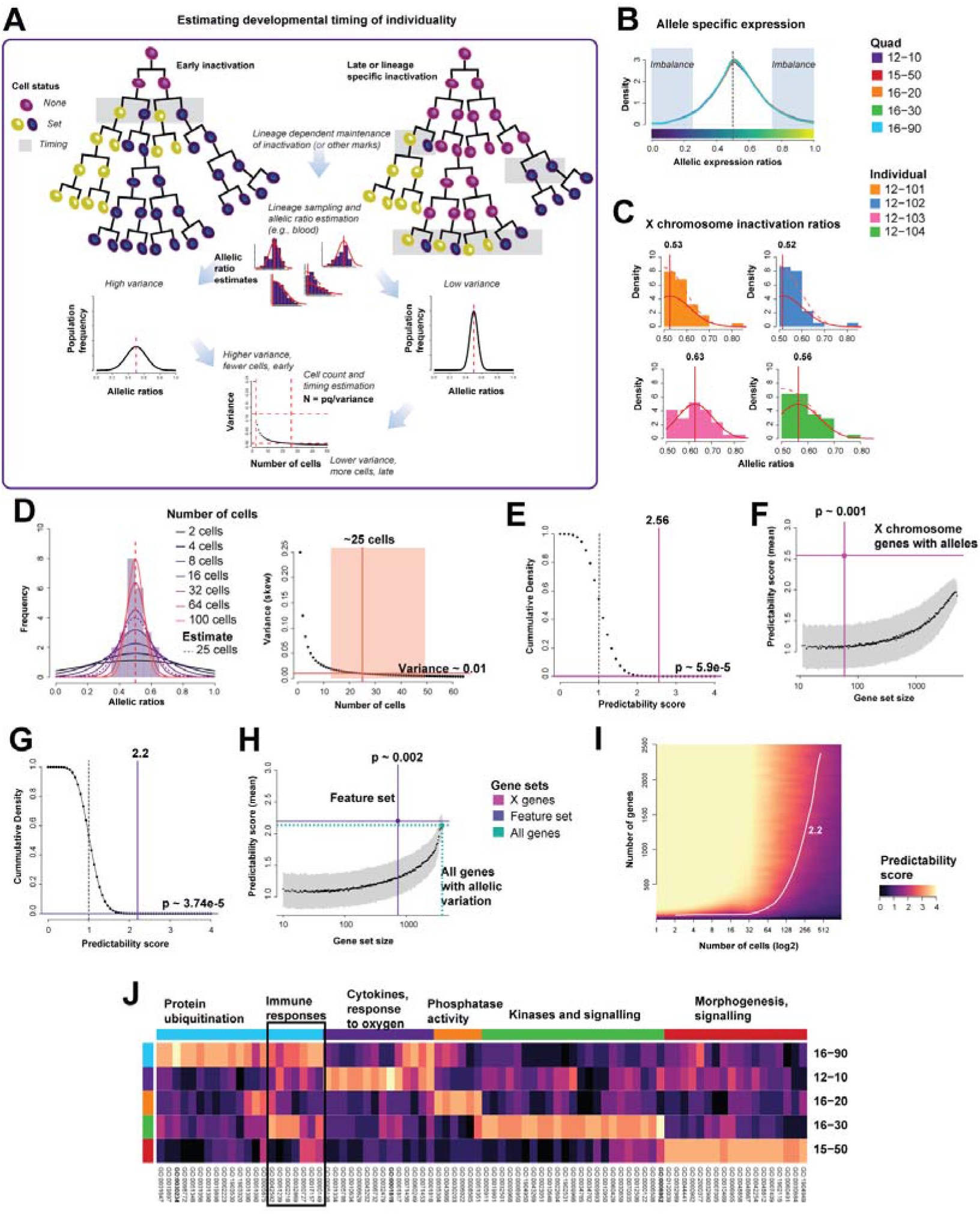
Persistent allelic imbalance as a mark of individuality. (A) Developmental timing of epigenetic marks can be estimated by calculating the starting number of cells required to generate the range of allelic imbalances observed. (B) Allele specific expression across all armadillos and all SNPs. (C) X-inactivation estimates from RNA-seq data in quadruplet 12-10 shows variation in skewing. (D) We estimate the number of cells where inactivation occurred to be around 25. (E-F) XCI is strongly predictive of identity. (G-H) Other genes with allelic imbalances also map identity. (I) Modeling imbalance, we find that our signal is explained by 500-700 genes and time events to a few hundred cells present when epigenetic marks were set. (J) And functionally this is explained by signaling and molecular functions unique to each quad.

More generally, we can define gene ‘identity signatures’ by their capacity to consistently identify a given individual, much like a fingerprint marks identity. This is a task that can be easily made into a formal assessment as a supervised classification problem and rigorously tested within our data (**Fig S9**). To this end, we developed a machine-learning method that determines the consistency over time of allelic profiles that distinguish armadillo siblings. In brief, we perform cross-validation, holding out one time-point as test data and learning allelic ratios that are distinct to each armadillo in the remaining training data. We then predict identity in the test data, reporting a score between 0 and 4, indicating how many individuals were correctly identified within the quadruplet sibling set (then averaged across all sets). Not surprisingly, the allelic imbalance ratios of the X genes are highly predictive of an individual within a quadruplet (**Fig. 2E**, score=2.56, p=5.9e-5, **Fig. 2F** p =0.001).

While an allelic signature is expected on the X-chromosome, we wished to extend this to a genome-wide assessment. Typically, allelic imbalance is attributed to the impact of a variant: either the variant within the gene has an effect on the stability of the mRNA, or an upstream SNV has cis-regulatory effects (*9*). As armadillos are genetically identical, any variant that has the same impact on gene expression will not distinguish individuals. Although we cannot exclude *de novo* variation, the expectation of the rate precludes it as the major factor. Rather, epigenetic regulation, set independently of genotype, will appear as random allelic imbalances, showing a preference for one allele over the other on average. For these imbalances to characterize individuals over time, they will need to be stably passed down along cell lineages. In essence, total imbalances will reflect the compositional distribution of these cell-lineages within an individual, where groups of cells are skewed in one direction, for a set of genes. Looking across all quadruplets and removing the genes on the X-chromosome, we find on average 700 genes within each quad to show an allelic imbalance, highly predictive of individuality (**Fig. 2G** score=2.2, **Fig. 2H** p=0.002) and implying a significant fraction of these genes are epigenetically canalized. In magnitude of impact, this provides a signature of individuality approximately equivalent to an additional half an X-chromosome worth of imbalance distributed across the genome of both females and males.

As in the case of XCI, we can estimate the timing at which expression was canalized by exploiting the variance of allelic expression under the same coin-flip (binomial) model. We find that most of the observed variance is due to epigenetic marks set early in the development of the blood lineage, at around a few hundred cells (**Fig. 2I**). Given the nature of epigenetic regulation, this is plausible, as the reprogramming of the embryo and setting of marks occurs at these stages (*10*). The signature of the allelic imbalanced genes was unique to each sibling cohort of quadruplets, but did include enrichment for a common immune response signal (**Fig. 2J**) as well as signals related to signaling and enzymatic activities. These are of particular interest in development as they ensure the switching on and off of programs that may result in phenotypic abnormalities if not controlled.

We applied the same paradigm to gene expression, as opposed to allelic ratios, to ascertain likely functional impacts of transcriptional individuality. Allele-specific expression is a useful probe for the timing of epigenetic effects, but does not directly assay the overall impact of total abundance reflected by a gene’s expression. As in the previous analysis, we define genes as markers of identity if their expression level can be used to predict identity at a held-out time point when trained on the remaining data (**Fig. 3A**). Consistent with greater selection for stability in gene expression, we find a weaker although still statistically significant signature of individuality (**Fig. 3B**, score=1.8, **Fig. 3C** p=0.014). Interestingly, allelically imbalanced genes do not show excess variability across individuals in their total expression – meaning these genes were stably expressed even if their alleles were not (**Fig S14**). This supports the hypothesis that selection is operating principally on expression level rather than at the level of individual alleles.

**Fig. 3:**
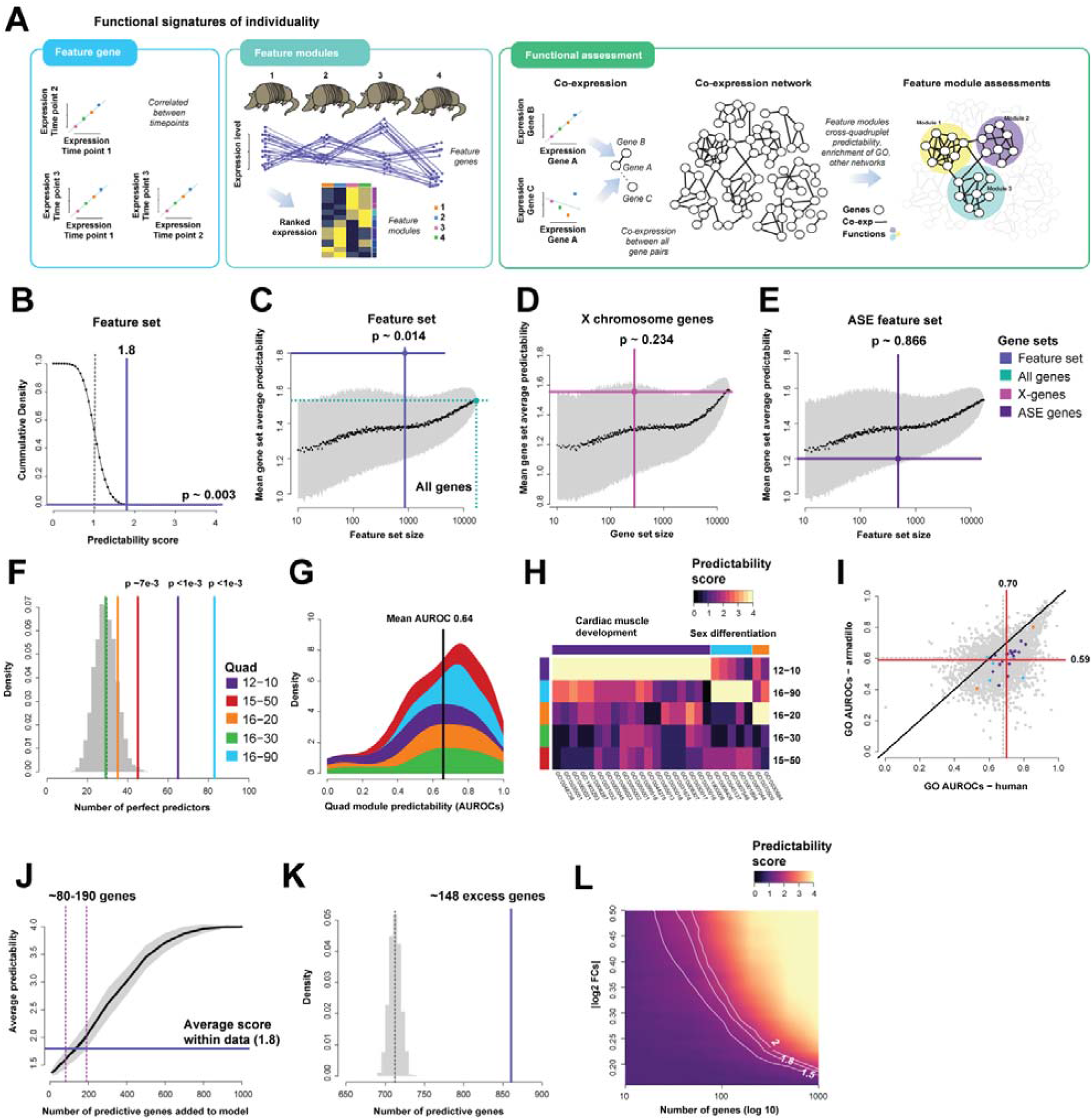
Gene expression marks functional signatures of individuality. (A) Feature genes mark identity and form feature modules (co-linear/co-expressed). (B– C) Feature gene sets mark identity, and this signal differs from the (D) X-chromosome genes and (E) ASE imbalanced genes since these sets are not predictive. (F) The signal is specific to each quadruplet (G) with weak cross-quad predictability. (H) These identity genes contribute to known pathways and process that are also quadruplet specific. (I) Globally, gene sets retain functional similarity across species, as shown by the average GO performances of their co-expression networks. (J-K) A model and our empirical estimates ~150 or so genes would contribute to this signal. (L) This reflects ~ 0.3 |log2FC| according to our model.

Are particular molecular functions more likely to vary across individuals? To answer this, we use a combination of co-expression and enrichment to identify and annotate molecular functions. Importantly, we find an uneven impact in the signatures of individuality across sibling cohorts, with individuality easier to detect in some quadruplet sets than others. Two of the armadillo quadruplets have many more genes which perfectly mark identity than expected (**Fig. 3F**). We define these gene sets as “perfect predictors”. Remarkably, the perfect predictor sets of genes are co-expressed in all armadillo quadruplets, even where they are not markers for identity. Because co-expression often reflects a shared functional relationship, it appears that the perfect predictors reflect pathways that, while present in all armadillos, were only differentially regulated in a few of the quadruplet sets. These genes are part of conserved functions that interact with environmental or lifestyle stimuli, such as sexual characteristics (hormones, heat) or cardiac muscle growth (exercise, stress). The perfect predictor genes are not outliers or distinctive in terms of their average expression levels (**Fig S12**). We estimate that individuality is encoded by a small fraction of the genome (**Fig. 3J,K**), with approximately 150 genes on average showing important alterations in gene expression level. Simulating the impact of this number of genes within our data suggests that relatively modest fold changes across this set of genes could account for the observed variation (|log2FCs|~0.3, **Fig. 3L**). In order to determine if the functional perturbations driving identity in armadillos were conserved in human, we assessed a large compendium of human expression data, finding broad similarity of co-expression across species, and suggesting the same functions are potential targets of developmental stochasticity in human as well (**Fig. 3I**).

While similar molecular functions are at play in humans and armadillos, we wished to attempt a more direct replication of our experiment in the human data best mimicking armadillo polyembrony, i.e., that taken from human identical twins. Identical twins share an *in utero* environment, but their post-natal environment is more variable compared to armadillos raised under controlled conditions (**Fig. 4A**). This is likely to make expression a less reliable measure of phenotypic individuality in humans: There will be additional variation reflecting uncontrolled lifestyle differences or environmental variation. And indeed, we see this reflected in the average fold changes in human monozygotic twins compared to armadillos. Performing pairwise expression comparisons between armadillos within a quadruplet set, we find the range of fold changes is very low (on average |log2FCs|~0.16, **Fig. 4B**). This is not surprising given the signatures of individuality we measured earlier – only a handful of genes will be differentially expressed. In contrast, human twins on average have fold changes closer to |log2FCs|~0.38, greater than a 1.2-fold average difference. We interpret this as arising due to uncontrolled environmental variation, as the average variation across unrelated armadillos in a fixed environment is also lower (armadillos |log2FCs|~0.30, humans |log2FCs|~0.56). Interestingly, genes showing variability between identical armadillos and humans are frequently perturbed in differential expression studies (**Fig. 4D**). This implies that the variation in gene expression is regulated, and that these genes are stimulus-responsive. Additionally, we find that these genes are less likely to be associated with eQTLs (**Fig. 4E**), indicating that their contribution to phenotypic variance does not arise through genetic variation, even outside of identical siblings. In summary, our data suggests that the regulatory factors at play early on during reprogramming define baseline phenotypic states which may vary stochastically between genetically identical individuals, but that a significant fraction of expression variation in humans reflects transient environmental responses, revealed as much larger fluctuations in expression levels (**Fig. 4C**).

**Fig. 4:**
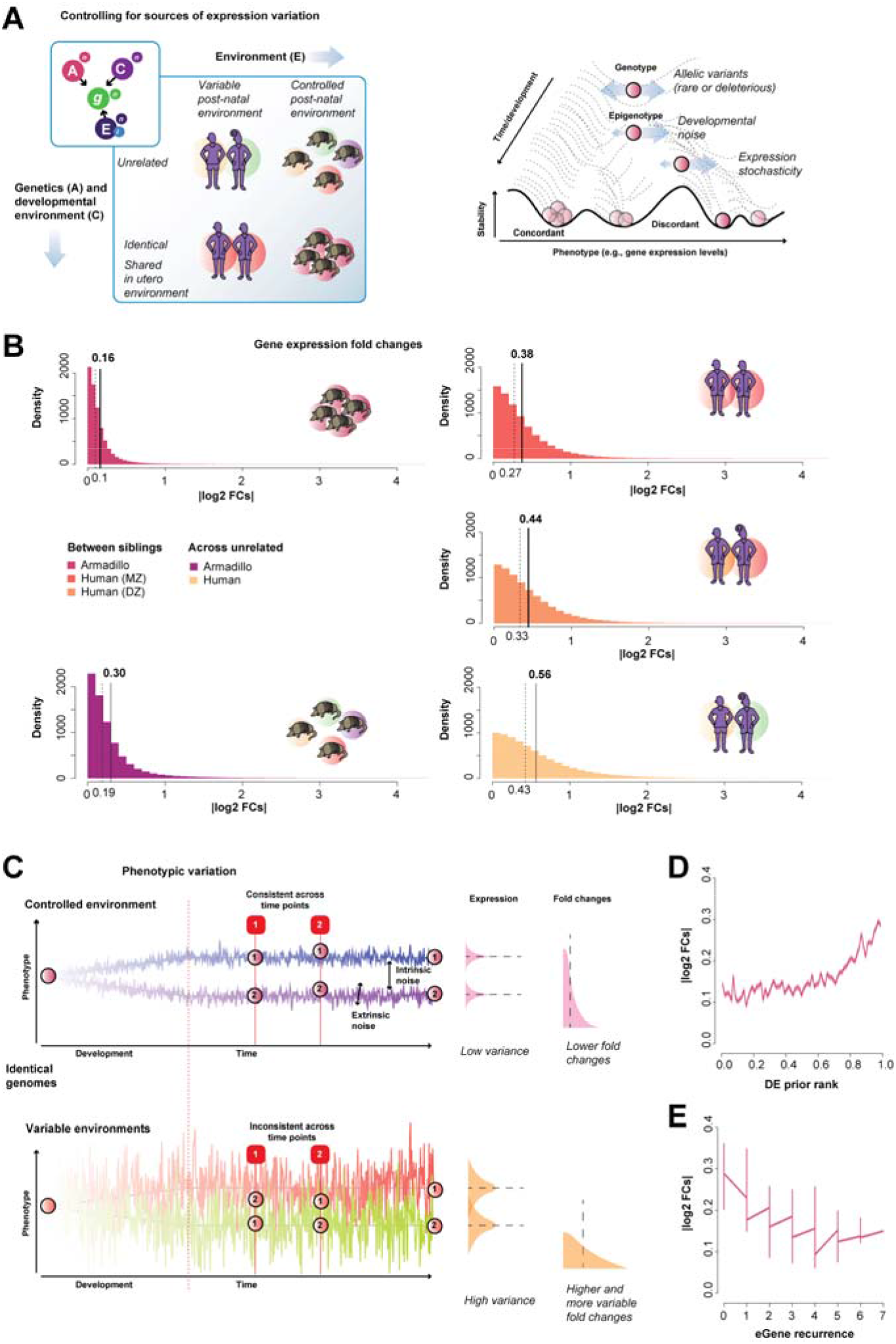
Armadillos reveal sources of variation. (A) Phenotypic variation results from three main sources: genetic variation, epigenetic variation and environmental effects. These are all influenced by stochasticity of gene expression. (B) Distributions of |log2FCs| calculated between armadillos within the same quadruplet, and those across quadruplets. Shown in comparison to fold changes between human twins (MZ and DZ), and those between unrelated individuals. (C) Modeling signatures of identity as observed in controlled and variable environments (D) Genes variable between identical siblings are commonly differentially expressed in the general population. (E) eGenes (genes with known cis-eQTLs from blood) show a trend towards lower fold change within quads, suggesting variability in the population is not genetically influenced.

In conclusion, our work demonstrates that purely stochastic variation in development has a large and permanent impact on gene expression, permitting the identification and characterization of genetically identical individuals over time. Using allelic imbalances, we have timed the contribution of developmental stochasticity within our data to the assignment of tissue-specific epigenetic marks. Using expression profiles, including co-expression and human twin data, we have determined conserved functions affected by developmental stochasticity. Given our cohort size, it is harder to estimate the magnitude of this effect downstream on phenotype; however, it is striking that in one of our two highest performing quadruplet sets, the impact on phenotype appears clear. The quadruplet set with high individuality in genes associated with muscle growth and development exhibits far more variance between siblings in size than the other quadruplets; in fact, the variability is roughly on par with the variance between quadruplet sets. As a Fermi estimate, we might imagine this suggests developmental stochasticity accounts for 20% as much variability as genetics does, in aggregate, for many phenotypes (e.g., perhaps 10% of total variance). Size is one of a handful of properties readily quantified, so we suspect this is simply a readout from one obvious phenotype exhibiting variability, rather than a discovery of the sole epigenetically variable phenotype. Because we did not select our cohort for disease, the influence of stochasticity on known variability in penetrance is likely far higher, consistent with the known effect of X-skewing. Through its potential influence on many phenotypes, developmental stochasticity defines a central convergent axis across individuals, species, and – probably – diseases. We believe, in time, that “noise” will cease to be a catch-all term and, instead, added to the traditional axes of nature and nurture as a principal and well-defined contributor to phenotypic variance.

## Acknowledgments

We thank Paul Pavlidis, Tony Zador, Peter Koo, Megan Crow and Dan Levy for helpful discussion and comments.

## Authors contributions

S.B. performed the analyses. S.B. and J.A.G. designed the study. F.M.K. collected the armadillos. L.B.A. and M.T.P. housed the armadillos and collected the blood. S.B. and J.A.G. wrote the paper.

## Data and materials availability

All data is available in the main text or the supplementary materials. The accession number for the sequencing datasets reported in this paper have been deposited in GEO under XXX and SRA under PRJNA591897.

## Material and methods

### Armadillo collection and samples

Five sets of armadillo quadruplets (20 armadillos in total) were used in this study. Pregnant females were captured using long-handled nets at night from the wild in 2012, 2015 and 2016. Capture of the pregnant females was done during the spring to avoid collecting females who were nursing young, but were potentially pregnant. The animals were retrieved from the nets and placed in kennels for immediate transport to the holding facility at the University of the Ozarks, Clarksville, AR. The pregnant females were kept in outdoor pens that had burrows where they gave birth to the quadruplets. The babies were kept with the mothers until they were observed foraging at about 6-10 weeks postnatal age. After separation from the mothers, the animals were housed together in semi-outdoors pens (rubber covered concrete floor under a roof). Most litters in the semi-outdoor pens shared the pen with another litter (2 litters per pen). All adults and young over 49 days postnatal age (pna) were fed a mix of dry dog and cat chicken-and-rice chow moistened with water—in an approximate ratio by volume of 1:1:2. Adults were provided 0.75-1.5 cups (indoor-outdoor) of moistened chow once a day during the gestation and 1.25-1.75 cups per day during known or suspected lactation. Animals housed in outdoor enclosures were able to forage as well. Occasionally, a raw egg and earthworms were provided in addition to the chow. Litters were fed replacement formula of reconstituted Esbilac puppy replacement formula until old enough (*11*). After 35 days pna, the diet was gradually transitioned to that of adult by 49-56 days pna. The wild-caught females were administered 0.15 ml Ivermectin SC, and Exceed antibiotic if they showed any wounds or abscesses. Adults were dewormed every 6-8 weeks with Panacure on chow for three consecutive days, or with 0.2 ml Ivermectin on food. The babies were treated once with Panacure on chow for three consecutive days.

At four to five months of age, the animals were delivered to the National Hansen’s Disease Program (NHDP) facility in Baton Rouge, LA where they were placed in pairs in modified rabbit cages (*12*). They were fed the same dry food as that given at the Arkansas facility. After a period of adaption of approximately one year, the animals were treated with Penicillin (1.0 mL) and dewormed with Ivermectin (0.1 mL) and Praziquantel (0.4 mL). Prednisone (10mg/mL) was also given at this time.

### Armadillo time course analysis and Hansen’s disease

Blood samples were collected at three time points per quadruplet staggered over the course of a year, starting from March 2017 until August 2018 (Table S1). The pilot study consisted of two sets of quadruplets (12-10 and 15-50), and then later blood was obtained from quadruplets 16-20, 16-30 and 16-90. During the course of the year, the original two of the five sets of armadillo quadruplets were infected intravenously in the saphenous vein with 1×109 Mycobacterium leprae derived from athymic nude mice (*13*) – both after the first time point was collected. Blood was collected at different time points throughout the course of disease and at 18-24 months post-infection, the animals were humanely sacrificed when they developed heavy M. leprae dissemination with severe hypochromic microcytic anemia. The bacteria will locate in the bone marrow and the animals will eventually succumb to secondary complications of persistent bacteremia if not sacrificed (*14*).

### Human twin samples

Blood samples from the TwinsUK data was used for the human twin analysis. Briefly, BAM files were downloaded from the EGAD00001001088 EUROBATS project after access was granted to the controlled files. This data contains 391 blood samples collected from monozygotic (MZ) and dizygotic (DZ) female twins. After quality control, we were left with 66 MZ and 96 DZ twin pairs available for analysis.

### Armadillo RNA-sequencing

Blood was collected from the subclavian vein in BD Vacutainer Glass Mononuclear Cell Preparation (CPT) tubes (Fisher, USA), and peripheral blood mononuclear cells (PBMC) were isolated following standard protocols (*15*). Blood collection was performed under general anesthesia using Ketamine HCL (10 mg/kg) and Dexdomitor (0.1 mg/kg). All animals were screened for leprosy and their health (CBC and blood chemistry) evaluated at tri-monthly blood screenings. RNA was extracted from the PBMC using an automated Maxwell 16 Instrument (Promega) and a Total RNA purification kit (Promega). Library preparation was done with a poly(A) selection kit (KAPA mRNA HyperPrep) to enrich for mRNAs. Multiplexed paired end sequencing (PE76) was done using an Illumina NextSeq500 on multiple flow cells. We blocked for lane batch effects by splitting the quadruplet samples into pairs and ran two pairs of each set per flow cell. We downloaded the armadillo genome (DasNov3) from Ensembl (v95) (*16*), and generated an index file for use within STAR (*17*). We mapped reads with STAR and standardized counts to counts per million (CPM) by summing the counts and dividing by 1e6.

### Armadillo DNA-sequencing

DNA was extracted from blood collected according to standard protocols. We sequenced each quadruplet together to obtain their identical genome sequence. We pooled DNA from all four individuals of a quadruplet, except in the case of quadruplet 16-30 where we could not get enough DNA from individual 16-301. An average of 2.3μg of DNA per quadruplet were sent for whole genome sequencing at the New York Genome Centre (NYGC). Library preparation was Illumina TruSeq Nano DNA, 450bp. Sequencing was done on the NovaSeq with 2×150bp. Coverage depth was 30X. Reads filtered on quality and were aligned to the DasNov3.0 genome from NCBI using BWA (*18*). Variants were called from the BAM files using the GATK Unified Genotyper (*19*) following best practices for DNA variant calling (*20*).

### Armadillo personal genome generation

We used g2gtools (unpublished, https://github.com/churchill-lab/g2gtoolsv0.2.0) to generate a personal quadruplet genome for each quadruplet set. We first created VCI files of the SNPs and INDELs using the g2gtools vcf2vci with the -pass and -quality tags. This is an indexed version of the VCF file required by g2gtools. Homozygous (alternate) SNPs and INDELs that passed quality control were kept. SNPs were incorporated into reference genome FASTA file using the g2gtools patch command. INDELs were then incorporated into the patched genome with the g3gtools transform command. A chain file was generated using the g2gtools vcf2chain command. We updated the genome annotation file (liftover) using the new genome with the g2gtools convert command. As the genome of the armadillo is not assembled beyond a large number of scaffolds, the patches and transformations were done per scaffold. Once completed, we concatenated all the scaffold FASTA files back into one. With these five personal genomes, we generated individual STAR indices. Using samtools (v.1.9), we generated index files for the new genomes, and dictionary files with picard from GATK (v3.6.0) (*19*).

### Armadillo personal genome mapping and allele specific expression analysis

Following quality control, we mapped reads from each quadruplet to their personal genome with STAR (v2.7) (*17*). The resulting bam files were then run through GATK’s v3 best practices pipeline (*20*) to filter for quality alignments. Briefly, the pipeline involves adding read groups, marking duplicates, and then splitting and trimming based on CIGAR. A WIG file was then built using the count command in IGVTools (v2.3.80) (*21*). We then generated a VCF file with the heterozygous and homozygous (alternate) SNPs for each quadruplet. This VCF file was converted to a BED file, and then liftover to update the coordinates to the personal genome. This was then converted back to a VCF file. The SNPs (VCF) and counts (WIG) were then overlapped to obtain allele specific counts. Once again these were all performed on individual scaffolds, and recombined at the end of the analysis, which allowed for parallelization of the pipeline.

### Defining the armadillo X-chromosome

As the genome of the armadillo is unassembled, we constructed the X-chromosome by identifying which scaffolds were most syntenic to mammalian X-chromosomes. As the X-chromosome has high synteny between mammalian species (e.g., mouse and humans 95%), we used alignments of armadillo scaffolds to the X-chromosome of both human and mouse. We used the UCSC (*22*) chain/liftover files between the armadillo genome and the human (hg38) and mouse (mm10) genomes. We extracted the scaffolds from these files that align to the respective Xs of the species. There were over a million human alignments (1,231,264) to around 2K armadillo scaffolds. The largest and most overlapping to the human X was scaffold JH573670.1, but holds no annotated human X homologs. To the mouse, there were less than a million alignments (873,607) to around 1.6K armadillo scaffolds. The largest is once again scaffold JH573670.1. We included smaller scaffolds with a high overlap (90% alignment) with the human and mouse X as the remaining potential X scaffolds. We consider these scaffolds to represent most of the X-chromosome of the armadillo. As a final X identifier, we located an *XIST* homolog which is not annotated in the current annotation. Using the human *XIST* sequence (NC_000023.11), we performed a BLAST (*23*) search on the armadillo genome. Of the 16 hits that were to annotated armadillo genes, we then performed a reverse BLAST on the human genome to find the reciprocal top hits. The two genes (ENSDNOG00000033080 and ENSDNOG00000047775) match to two *XIST* exons, and both these genes belong on the same armadillo scaffold (JH583104.1) and are within a few 100Kbp. These two genes were also hits using the mouse *Xist* (NC_000086.7). Using these genes as placeholders, we could derive the rest of *XIST* from the read pileups. The locus is JH583104.1:145,010-175,550.

### Building functional annotation sets for the armadillo

Currently, no gene functional annotations exist for the armadillo. We used the gene annotations from *Ensembl* (*16*) to generate a gene ID map between human and armadillo homologs. From the total of 33,374 coding and non-coding genes and transcripts annotated for the armadillo, there are 13,492 human homologs. In close parallel to the GO(*24*) annotation project’s own process, we built an armadillo ontology using human gene-GO annotations(*25*). Within our current mapping, on average, each armadillo gene belongs to ~85 GO groups, and each GO group has on average ~56 genes.

### Measuring transcriptional stochasticity

We performed a transcriptome wide cluster analysis of the samples using Spearman’s rank correlation as a measure of sample-sample similarity. To find highly variable genes (HVGs) within each quadruplet set, we performed variance analyses and ranked genes based on their coefficient of variation and mean expression levels. We measured similarity of these variable genes for recurrence across the quadruplet sets. Using the set of top 100 HVGs of a quadruplet sets at a given time point, we calculated their AUROCs in cross-quad comparisons. Additionally, we ran an ANOVA across all the samples of a quadruplet to identify genes that were most variable within a quadruplet.

### X-chromosome inactivation analysis and cell number estimates

For every female armadillo, we estimated the X inactivation ratios of genes with alleles. For this, we took the variants called on the X scaffolds. For each gene, we combined the three timepoints by adding the count data. Since we do not have phasing information but wished to summarize the allelic ratios to a single gene, we took the most powered SNP (that with the most reads) as the representative SNP and calculated the allelic ratio as that of the reference to total count. These gene ratios were then used to estimate the X skewing ratio for the individual female armadillos. There, we fitted a folded normal to the ratios and used the maximum log likelihood estimate to obtain the estimated overall skew. Finally, we took the variance of the extremal estimated skew values to estimate the number of cells (N) in the original starting pool(*26*). The formula for the variance of a binomial distribution was used. Since we assume that the probability of a cell inactivating either X is 0.5, p = q = 0.5 so the formula becomes:

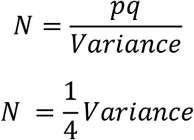

### Measuring identity

As a test for individuality, we developed a machine-learning method that tests for the relative consistency of expression across an armadillo quadruplet across time. This is equivalent to identifying differentially expressed genes, but rather than looking between two conditions or two individuals, it is across four. The idea here is that differentially expressed genes in this way are indicators of identity. We first select a feature set of genes based on correlations between two time points. For each gene, we calculate the Spearman rank correlation between the values across a quadruplet for one time point and a second time point. If the rank ordering is consistent (i.e., the correlation is 1), then this gene is selected as a feature gene. We then test for consistency in the third time point. As the first two time points are perfectly correlated, these genes form the training set, and the left out time point is the test set. A gene scoring matrix (4 by 4) is built per gene by comparing the ordering of the test and training data. Each individual gives a score of 1 to the test data individual it thinks it is (i.e., which rank it matches), and a 0 otherwise. We then sum all the feature gene scoring matrices to produce an aggregate scoring matrix. Then, in a winner takes all strategy, we calculate a score which represents the number of armadillos that correctly predict themselves. The final score is between 0 and 4, with 4 as perfect predictability i.e., each armadillo correctly identifies its future (or past) self. We repeat this three times, using the first and second time points as training, the first and third, and finally the second and third, and then testing in the left out time point. We average this across time points to get quadruplet specific scores, and also across all to get a final overall score for the analysis. We calculate an analytic p-value for this score by convolution of the expected distributions. We calculate an empirical p-value by repeating the learning task on randomly selected genes.

### Co-expression network analysis and functional enrichment using machine-learning

For each quadruplet at each time point, we constructed a co-expression network(*27*). Briefly, we calculated the spearman correlation coefficient for each gene-pair across the four individuals within a quadruplet. These values are used as weights for a gene-gene co-expression network, which is then rank standardized. The individual networks (a total of 15) were then summed to generate a final aggregate co-expression network. For all networks, we first applied unsupervised clustering analyses and tested for functional enrichment of the network modules identified by using our supervised machine learning tool (EGAD(*28*)) on the aggregate network and the individual networks. In EGAD, we report performance using AUROCs, as is typical within machine learning. An AUROC is the area under the receiver operator characteristic curve, and is the probability that a positive result is ranked higher than negative result by the algorithm (e.g., does this list of DE genes rank genes involved with “function X” higher than other genes). We used the functional annotations we mapped from GO for this assessment. As a test for cross-quad predictability, we aggregated the individual networks leaving one quadruplet out (LOQO). Then, with the modules identified as predictive of identity for each quadruplet, we measured the performance using EGAD within the aggregate with that quadruplet left out, as per typical cross-validation.

### Empirical models to estimate genes and lineage

From the identity analysis, we estimate the number of genes that could drive the signal through a series of empirical models. In our first model, we simulate our allelic identity experiment. We took the underlying allelic expression data and added a proportion of variance to a fraction of the genes. We then calculated the average identity performance. We then convert the variance of the underlying data to a number of cells estimate by extending the analysis from the X inactivation estimate. To summarize, we assume that autosomal genes that display allelic imbalances are under regulatory control and are being expressed either monoallelically (some cells express one or the other), or differentially (one allele is expressed at a higher or lower amount). We also assume that this is persistent across time, such that once a cell is committed to expressing an allele, its lineage will continue to express this allele at a similar or equal amount. We also assume that this choice is random in a pool of cells at the same time. With these in mind, we can estimate the number of cells and the fraction of the genome that gives rise to the performance observed.

For the gene expression model, we simulate two potential models. In the first, we assume the signal is localized. There, we take the underlying expression data and add a perfect signal to a fraction of the genes. We then measure the average identity performance and estimate the number of genes necessary for our measured signal. In a second model, we assume the signal is distributed. As in the allelic model, we simulate the expression identity analysis by adding a *M* proportion of variance (signal) to *N* number of genes. We then calculated the average identity performance that this signal would generate. We then converted the fraction proportion of variance to a log2 fold change (|log2FC|).

### Co-expression relationships shared across species due to conserved stochasticity

We constructed an aggregate human network across all extant blood data available within recount2(*29*). The human data is across 60 experiments (a total of 3,174 samples). We use “experiment” to refer to an entire expression dataset, across all its samples. After constructing the aggregate network, we used EGAD (*28*) to measure the networks performance with GO (*24*). We compared the AUROC GO performances across the species.

### Fold changes to measure phenotypic variation

As a measure of transcriptional (and phenotypic) variation between samples, we calculated pairwise log2 fold changes (|log2FC|) of gene expression values. We normalized each sample to CPM, and then took the absolute value of log2 of the ratio of each genes CPM (*g*_*i*_ is a gene, *n* and *m* are individual samples). We calculated these for armadillos within a quadruplet and for armadillos across quadruplets. We repeated this with the monozygotic and dizygotic human twin data, and also pairwise across different pairs of individuals. To ensure that we are comparing similar genes, we restricted our analysis to approximately 7500 genes that were homologs and were in both expression data sets.

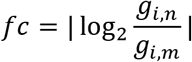

### Fold change comparisons to known transcriptional variation properties

A property of transcriptional variation of interest is the null/prior, i.e., genes commonly differentially expressed (DE). We wished to assess whether the genes fold changes are also influenced by cryptic genetic variation and compare this property (the DE prior rank(*30*)) to the fold changes calculated. We looked at fold changes due to non-genetic effects (i.e., the within quadruplet and human MZ twins). We also assessed the correlation of these DE prior properties to the fold changes that arise due to both genetic and non-genetic differences by comparing to cross-quad and cross-twin fold changes. Similarly, genes known to be influenced by eQTLs (eGenes) are of interest as they reflect transcriptional variation due to allelic differences. To this end, we collected 7 independent human studies (**Table S6**) and used the frequency of the eGenes across the studies as a measure of the likelihood that the gene is reproducibly an eGene. Once again, we compare these metrics to the fold changes for both human and armadillos.

