## Supplement for "The transcriptional legacy of developmental stochasticity"

#### Supplementary Figures

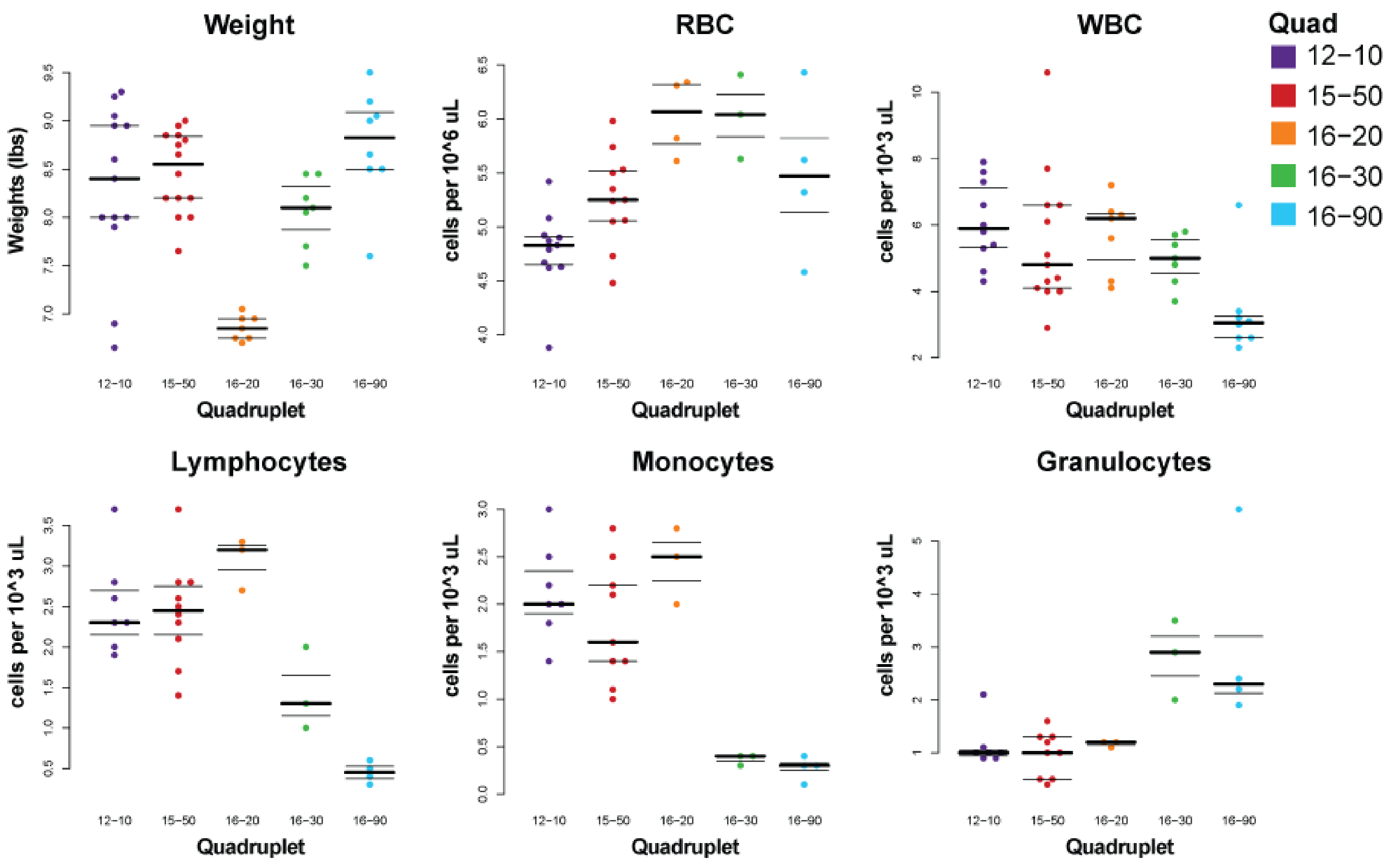

**Fig. S1 Weight and blood measurements of the armadillos.**
Multiple time points are included in the top row, showing consistency across time and within quadruplet. The bottom row shows cell counts for our third time point.

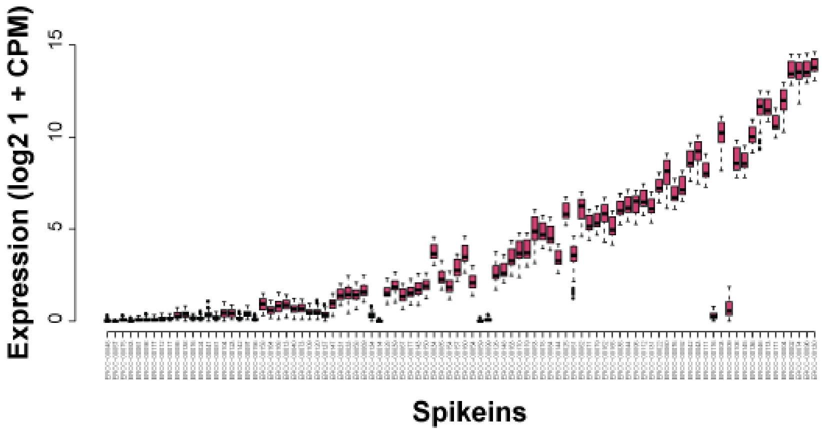

**Fig. S2 QC:**
Consistent expression levels for the spike-ins across the samples

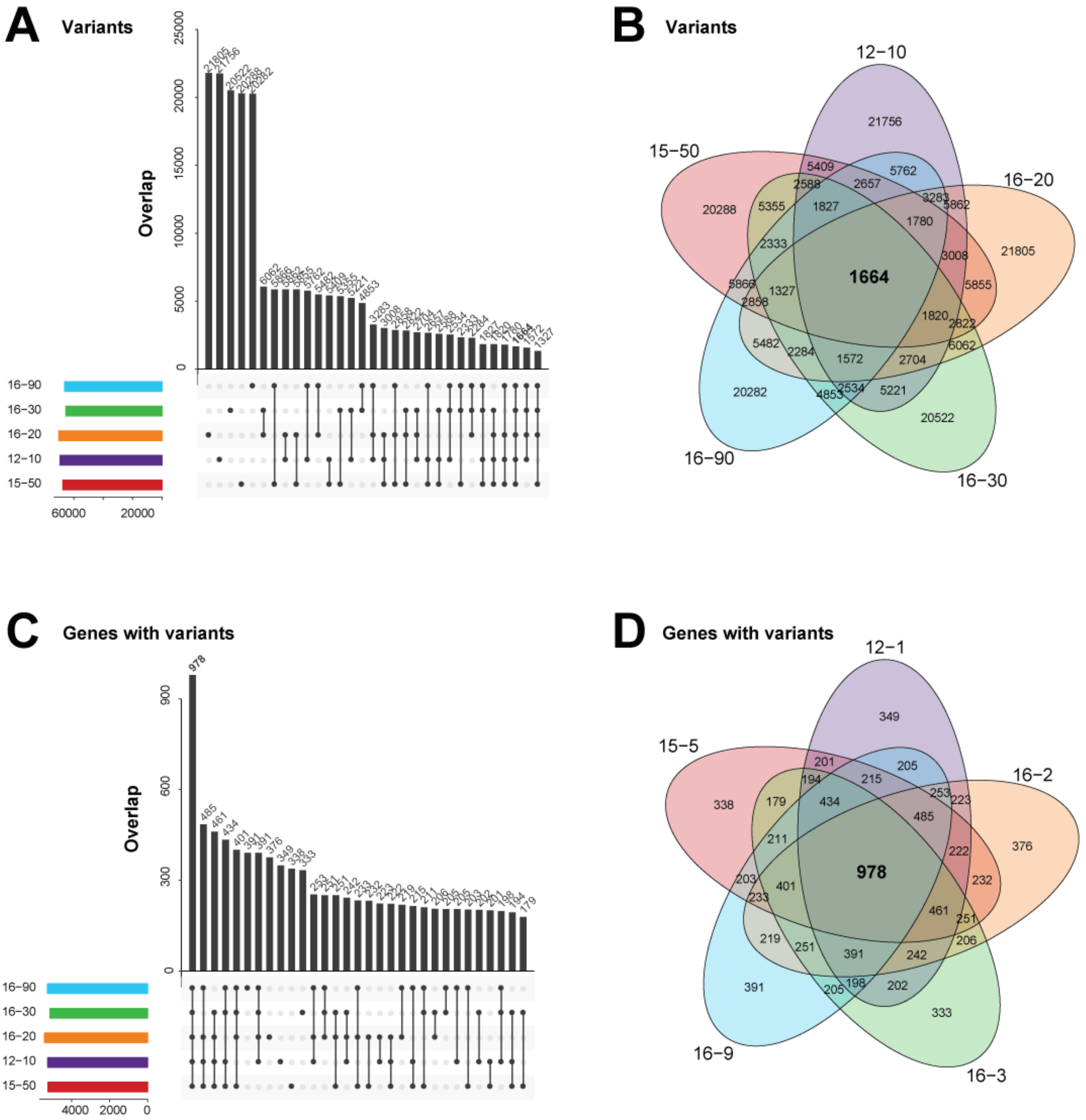

**Fig. S3 Genetic similarity across the quads:**
(A,B) The overlap of these variants called that are in genes are less than 2000 variants (out of a total of 70K per quad). (C,D) The variants fall in common genes (~900 out of ~5000).

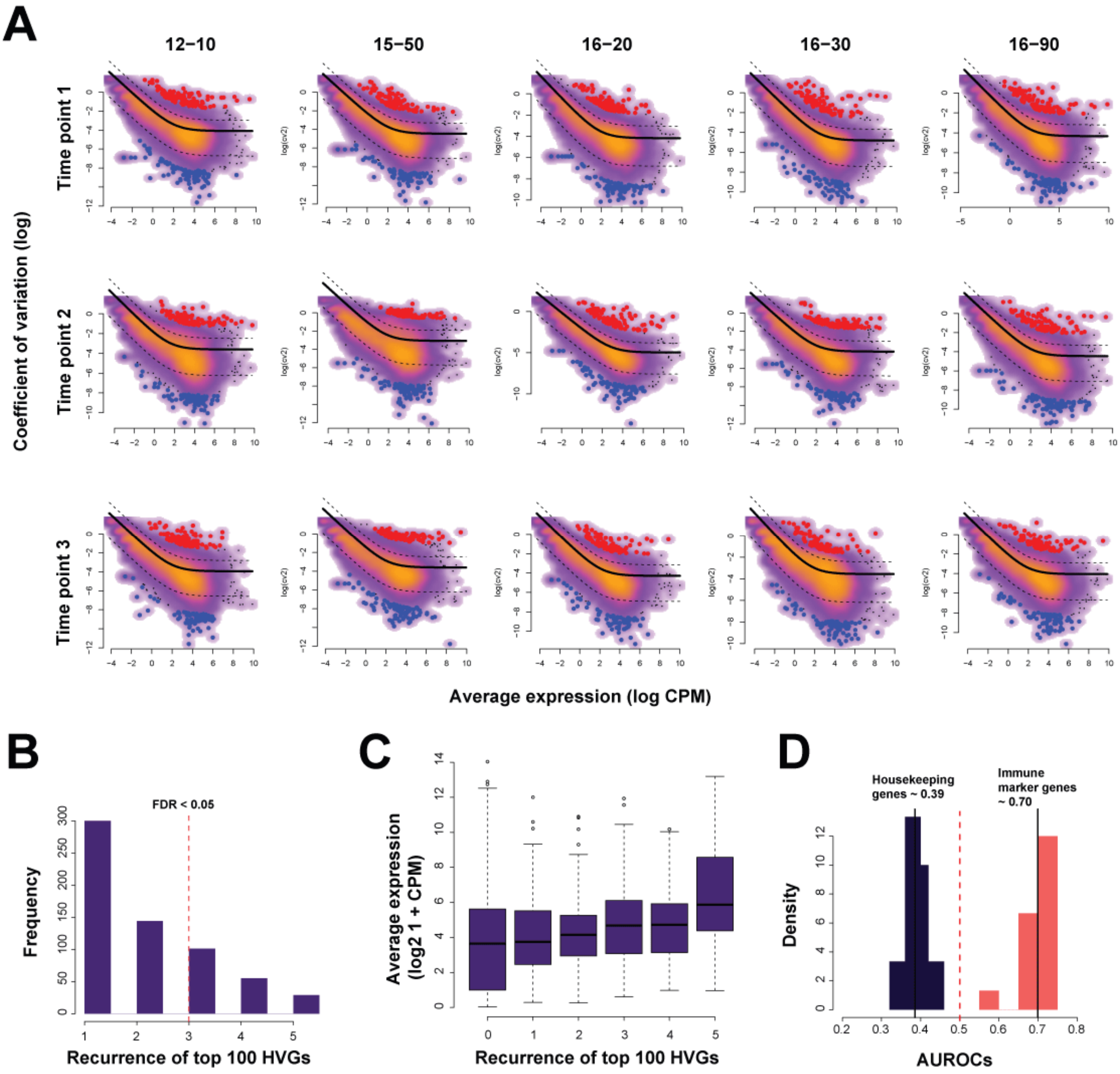

**Fig. S4 Variable genes are replicable across armadillos:**
(A) Average expression levels versus coefficient of variation was used to select the top 100 most variable genes (highlighted in red). (B) These genes are recurrent across the armadillos and therefore not very specific. (C) Additionally, the most recurrent variable genes also happen to be more highly expressed despite controlling for expression. (D) Overall, the genes are enriched for immune marker genes and depleted for housekeeping genes.

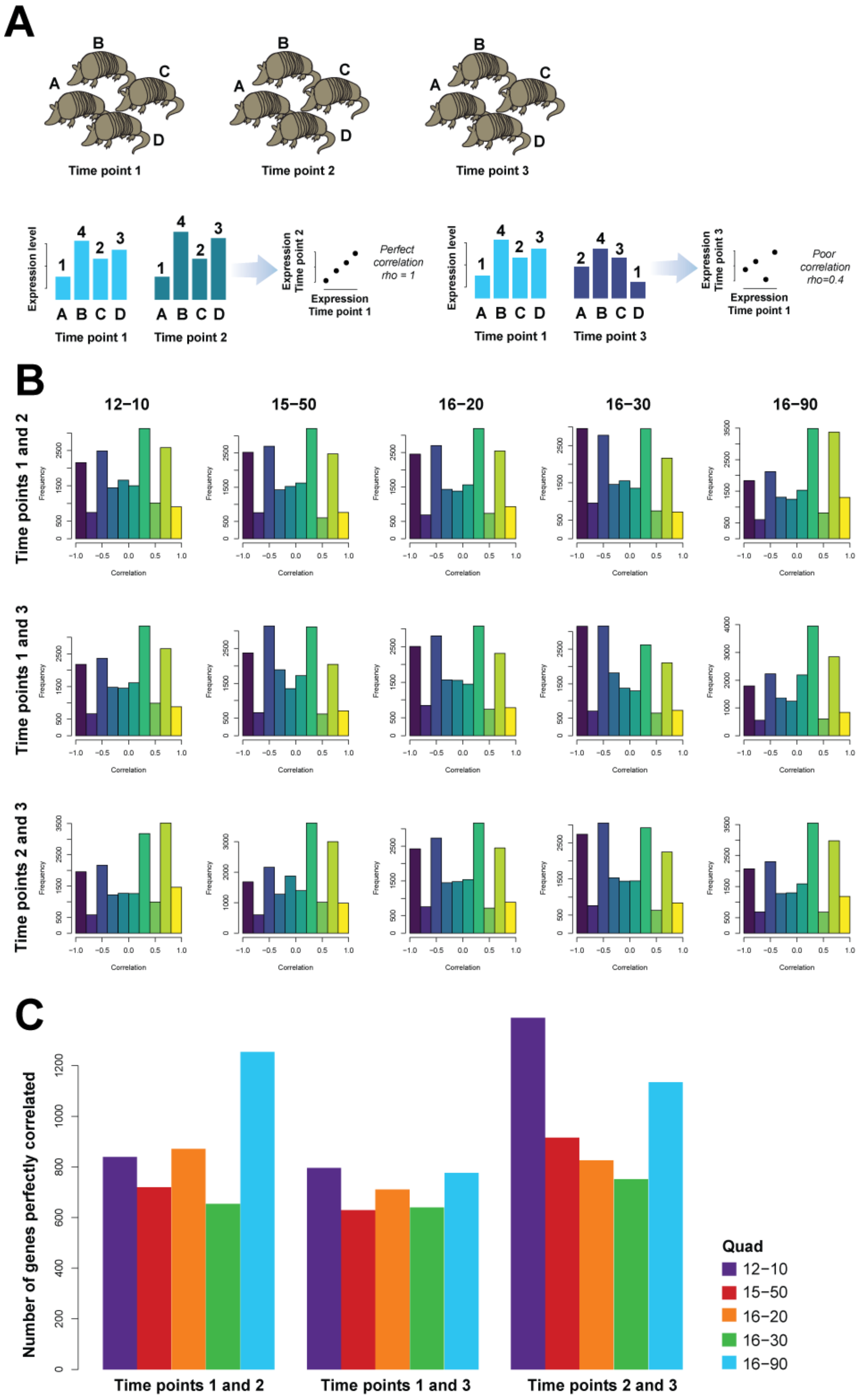

**Fig. S5 Genes correlated across time points.**

(A) For each quadruplet, we have three time points where we have sampled RNA. Each time point contributes a rank ordering of genes. We measure the consistency of this rank ordering by calculating the correlation between these orderings. Perfect correlations (rho = 1) are where the ordering is identical. Poorer correlations are where the ordering has shuffled. (B) Distribution of the correlations (per gene) across the individuals within a quadruplet between time points. (C) Tally of perfect gene correlations for each quad across time points.

**Fig. S1.**

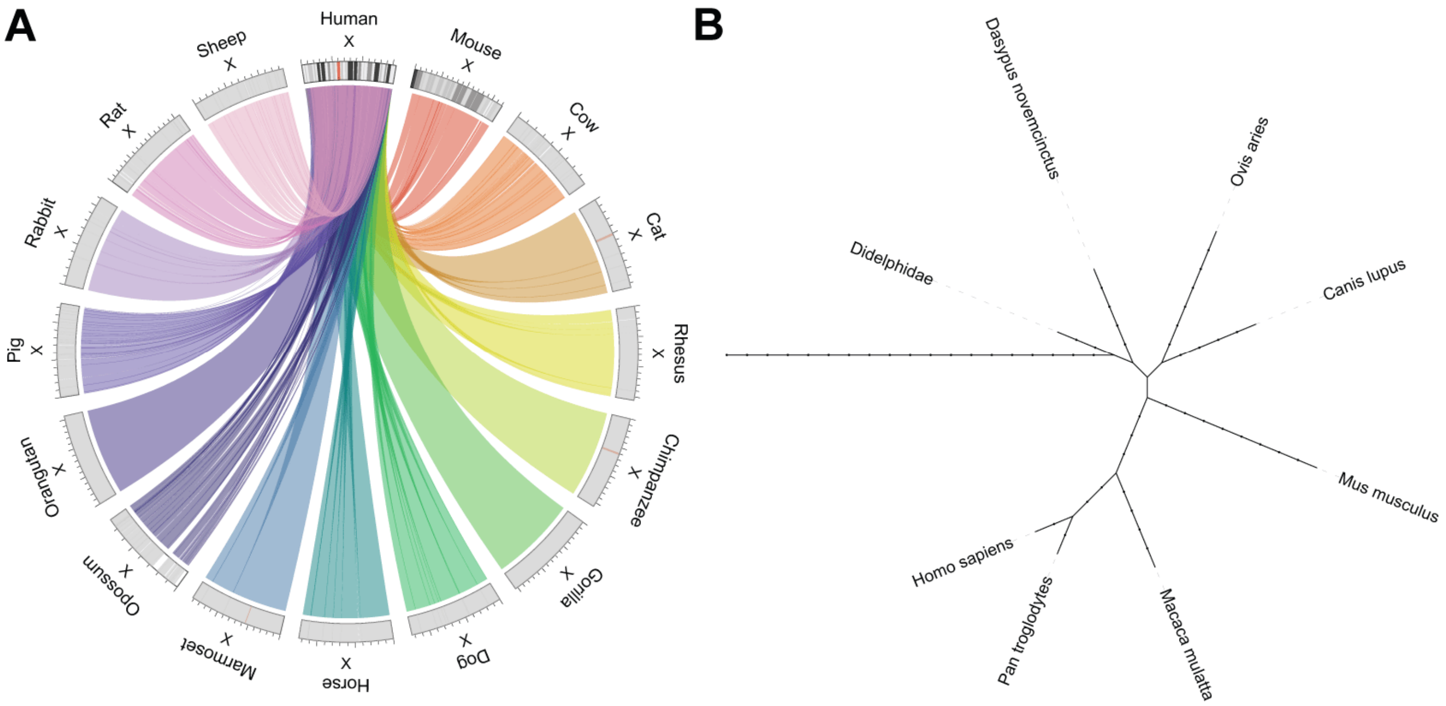

**Fig. S6 X-chromosome synteny:**

(A) Circos plot of X-chromosome synteny across annotated species. (B) Phylogenetic comparisons. Plots generated from: http://bioinfo.konkuk.ac.kr/synteny_portal/htdocs/synteny_circos.php and https://phylot.biobyte.de/index.cgi

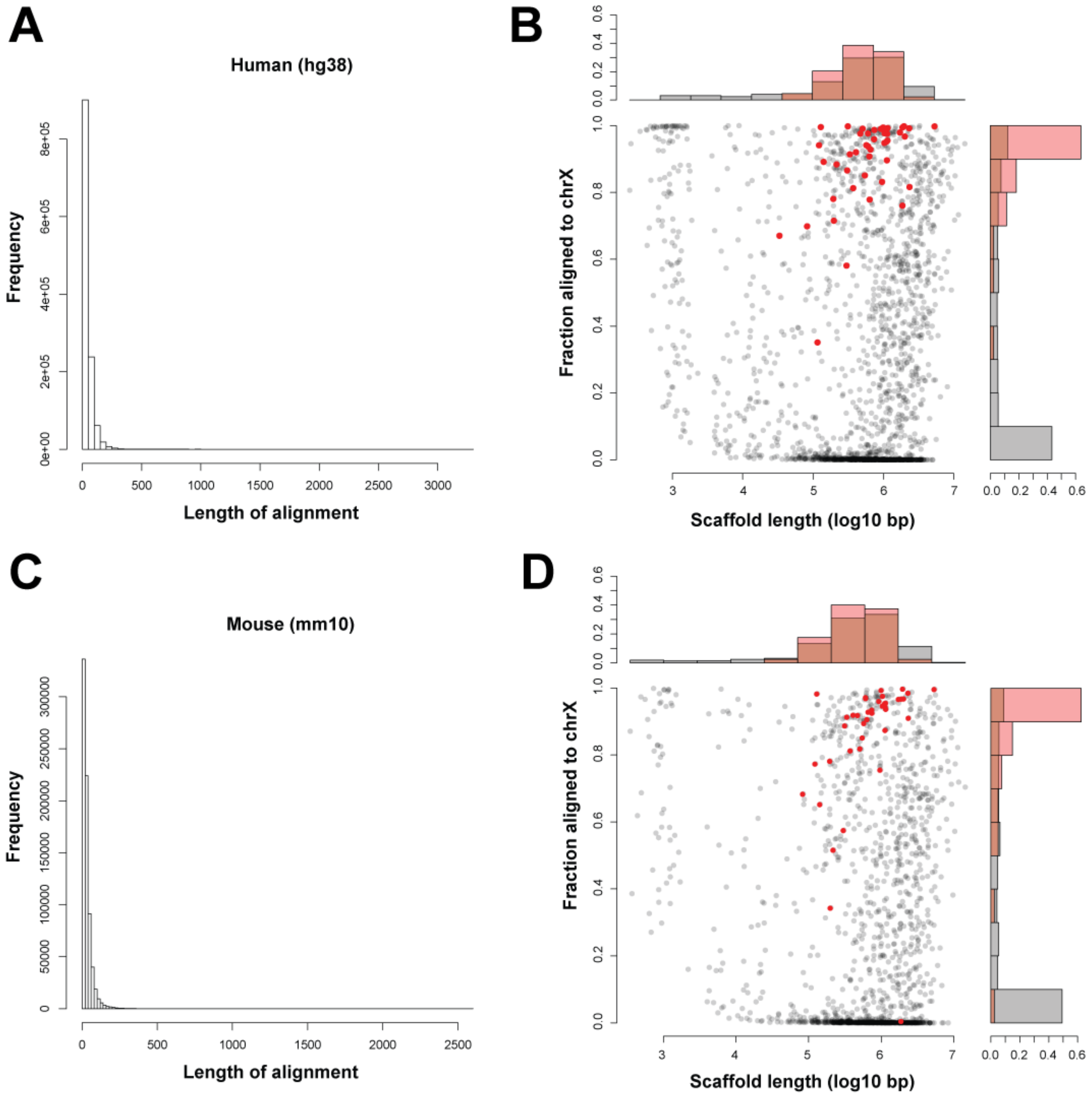

**Fig. S7 Scaffold length versus fraction aligned to X-chromosomes:**

(A) Distribution of alignment lengths of the human X chromosome to armadillo scaffolds. (B) Armadillo scaffold length versus proportion of scaffold that maps to the human X chromosome. (C) and (D) same as above but for mouse. Red points are human and mouse X-chromosome gene homologs (respectively).

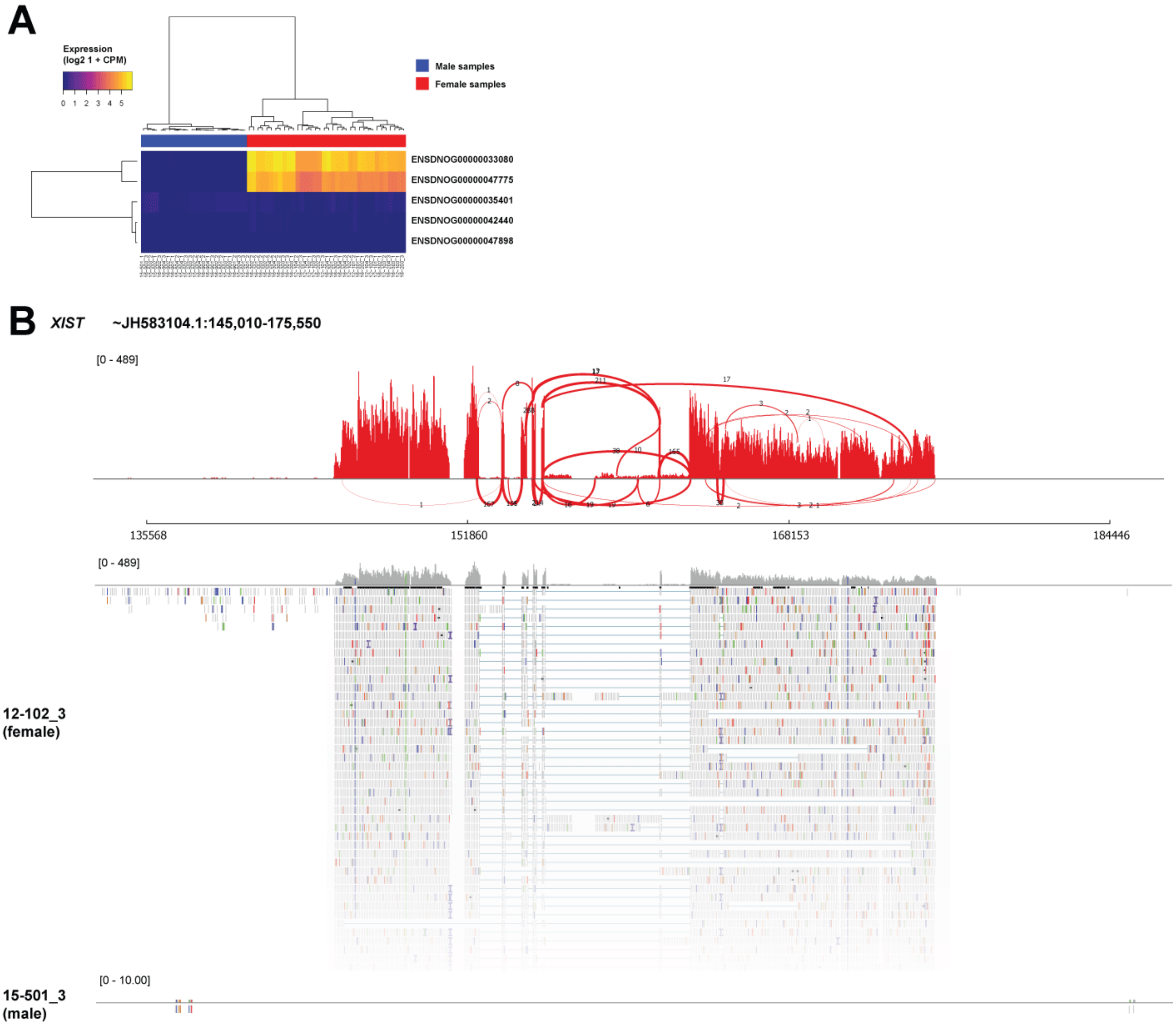

**Fig. S8 It eXISTs:**

(A) Expression of genes annotated near top BLAST hits. (B) Sashimi plot of predicted *XIST*.

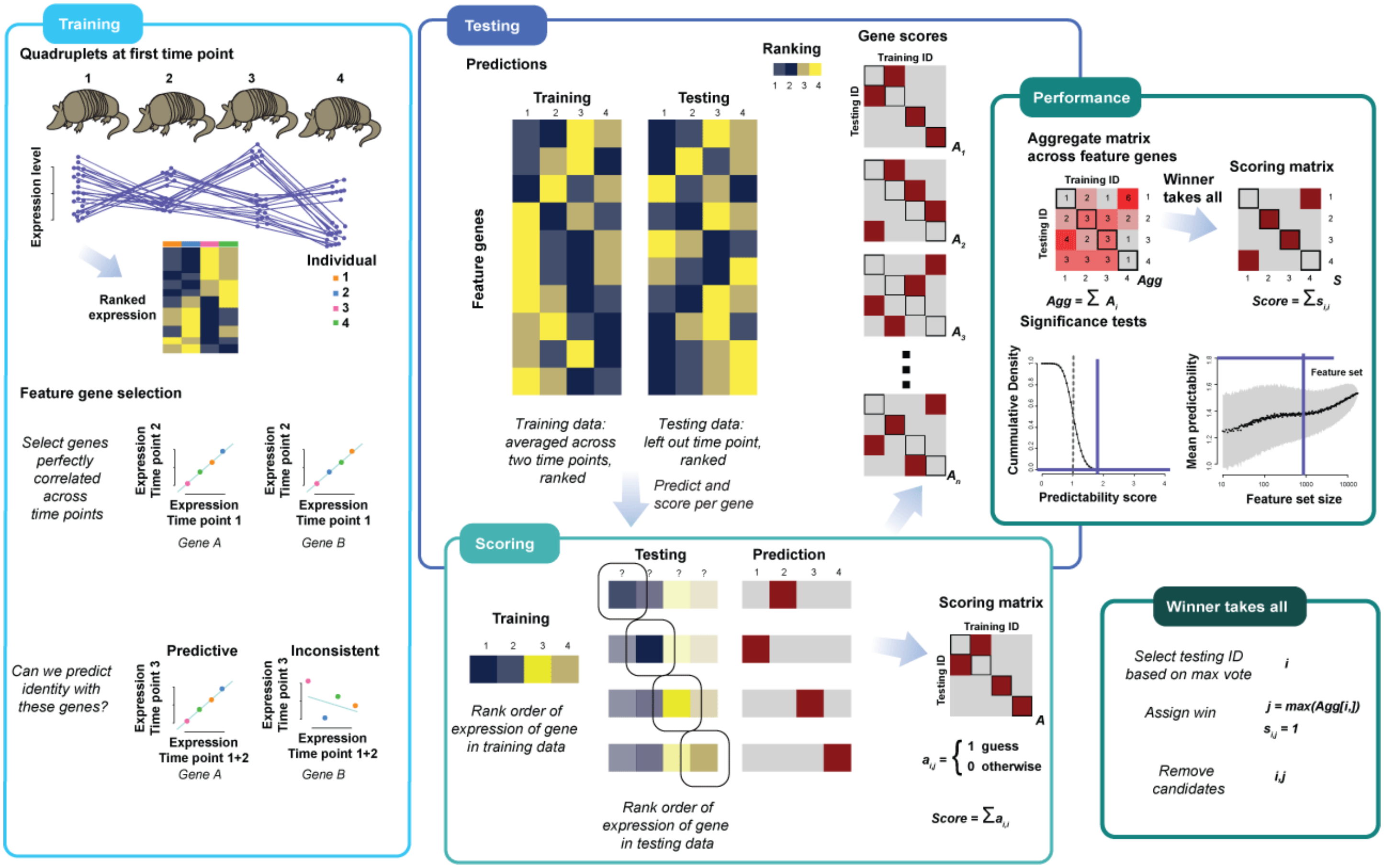

**Fig. S9 Testing for individuality:**

Schematic of machine-learning method. Training: feature set of genes are selected based on correlations between two time points. For each gene, we calculate the Spearman rank correlation between the values across a quadruplet for one time point and a second time point. If the rank ordering is consistent (i.e., the correlation is 1), then this gene is selected as a feature gene. Testing: The first two time points are perfectly correlated, these genes form the training set, and the left out time point is the test set. A gene scoring matrix (4 by 4) is built per gene by comparing the ordering of the test and training data. Each individual gives a score of 1 to the test data individual it thinks it is (i.e., which rank it matches), and a 0 otherwise. Scoring: We then sum all the feature gene scoring matrices to produce an aggregate scoring matrix. Then, in a winner takes all strategy, we calculate a score which represents the number of armadillos that correctly predict themselves. The final score is between 0 and 4, with 4 as perfect predictability i.e., each armadillo correctly identifies its future (or past) self. Performance: We repeat this three times, using the first and second time points as training, the first and third, and finally the second and third, and then testing in the left out time point. We average this across time points to get quadruplet specific scores, and also across all to get a final overall score for the analysis. We calculate an analytic p-value for this score by convolution of the expected distributions. We calculate an empirical p-value by repeating the learning task on randomly selected genes.

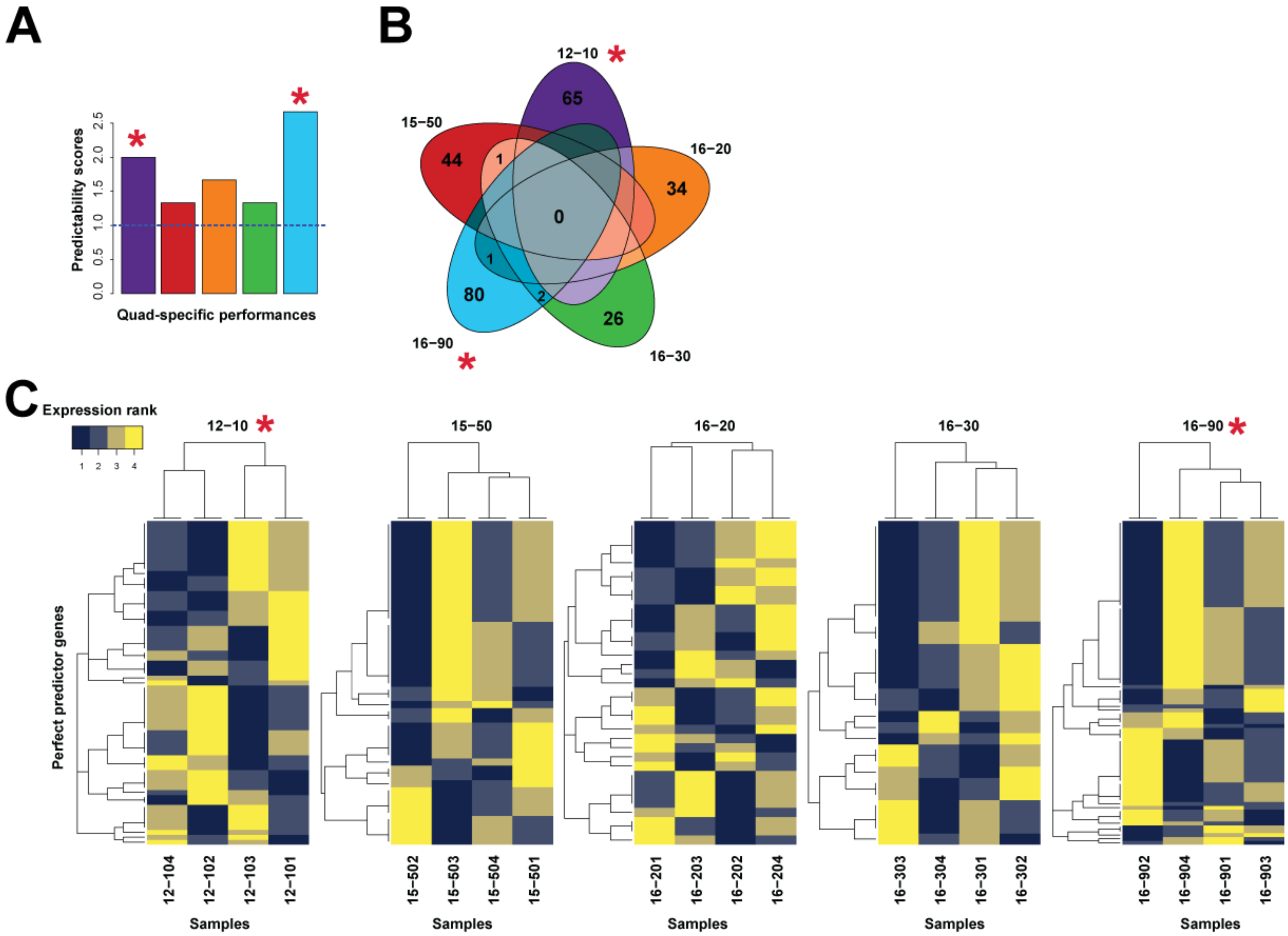

**Fig. S10 Perfect predictor genes and modules:**

(A) Average predictability performances of each quadruplet. Significant above the null are quads 12-10 and 16-60. (B) The number of perfect predictors (those predictive across all three time points) are low. (C) Perfect predictors form modules within the data.

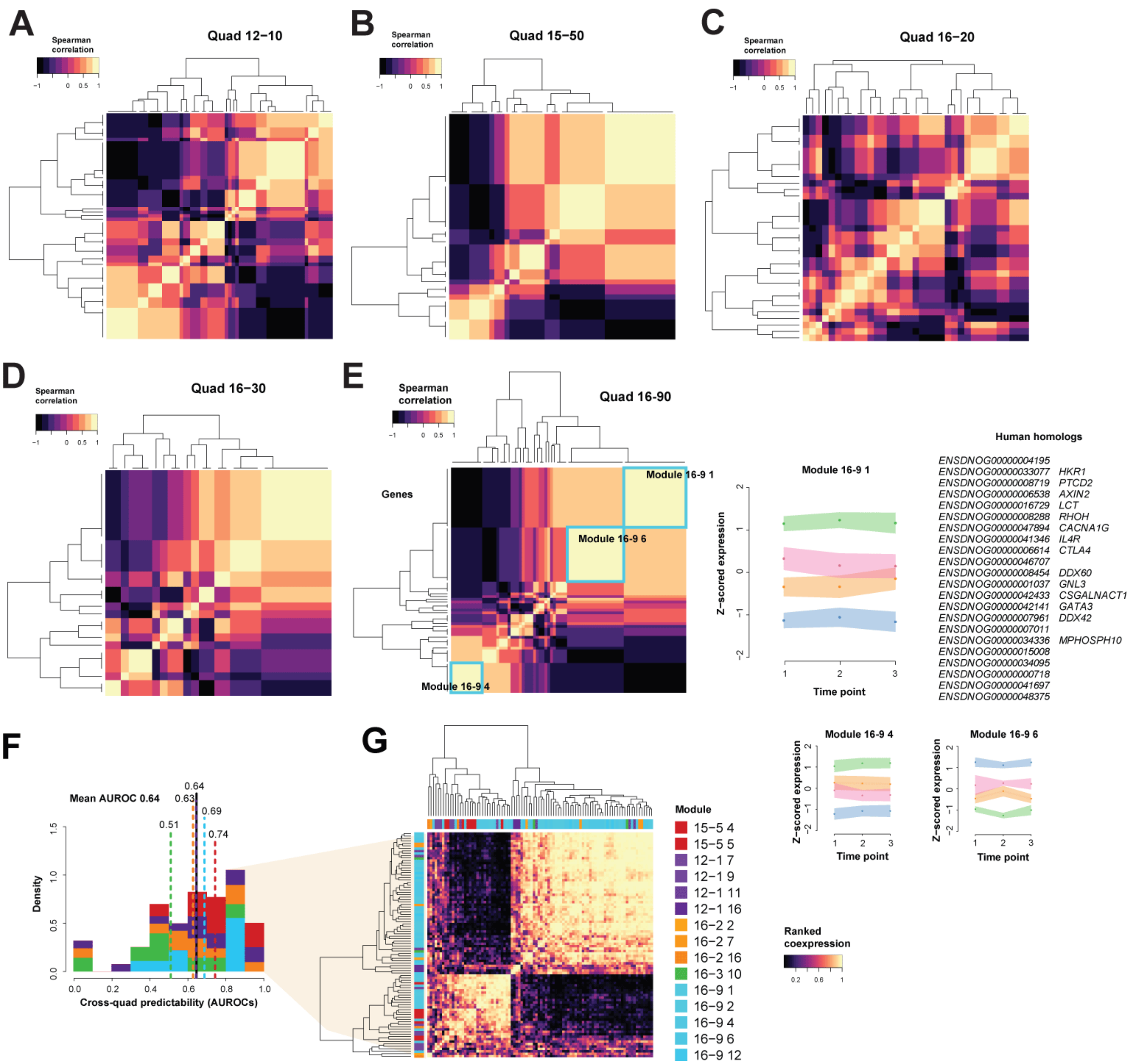

**Fig. S11 Perfect predictor genes and co-expression:**

Perfect predictors as co-expressed blocks for all quads (A-E) with quad 16-90's (E) larger modules highlighted.(F) Taking these modules and testing their performances using EGAD in aggregate co-expression networks. Each network is an aggregate of all networks generated excluding those of the quad's genes being tested. (G) The top performing modules and genes shown in the full aggregate network.

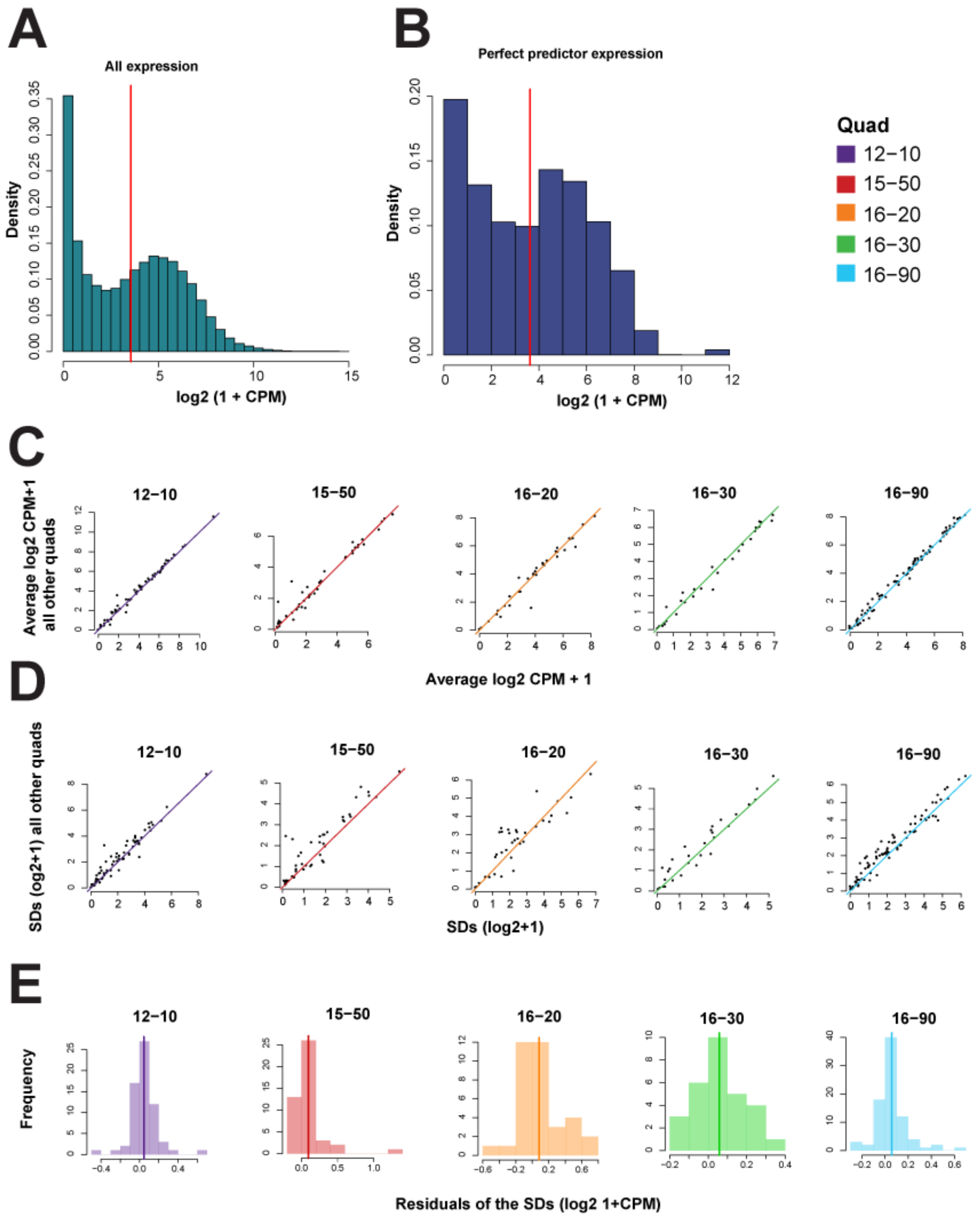

**Fig. S12 Characterization of perfect predictors showing normal expression properties:**

(A) Expression profiles of perfect predictor genes (B) are similar to background expression as shown in (A). (C) Scatterplots of average gene expression within a quad compared to all other quads. The expression values sit very closely along the diagonal. (D) SDs of perfect predictors compared across armadillos. (E) Distributions of residuals of variance within a quad compared to all other quads. If there is excess variance, we expect to see a positive shift. On average, no signal is apparent.

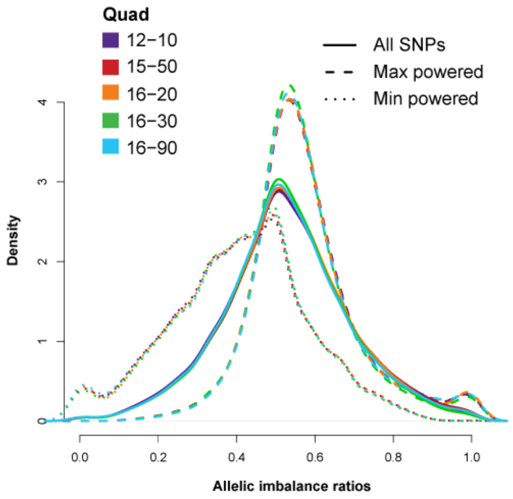

**Fig. S13 Allele specific expression imbalances:**

Some reference bias when taking the most powered SNP and alternate SNP bias when taking the lower expressed SNP. Thick lines are all imbalances. Large dashes are max-powered SNPs. Small dashes re min-powered SNPs. Min and max powered based on count numbers.

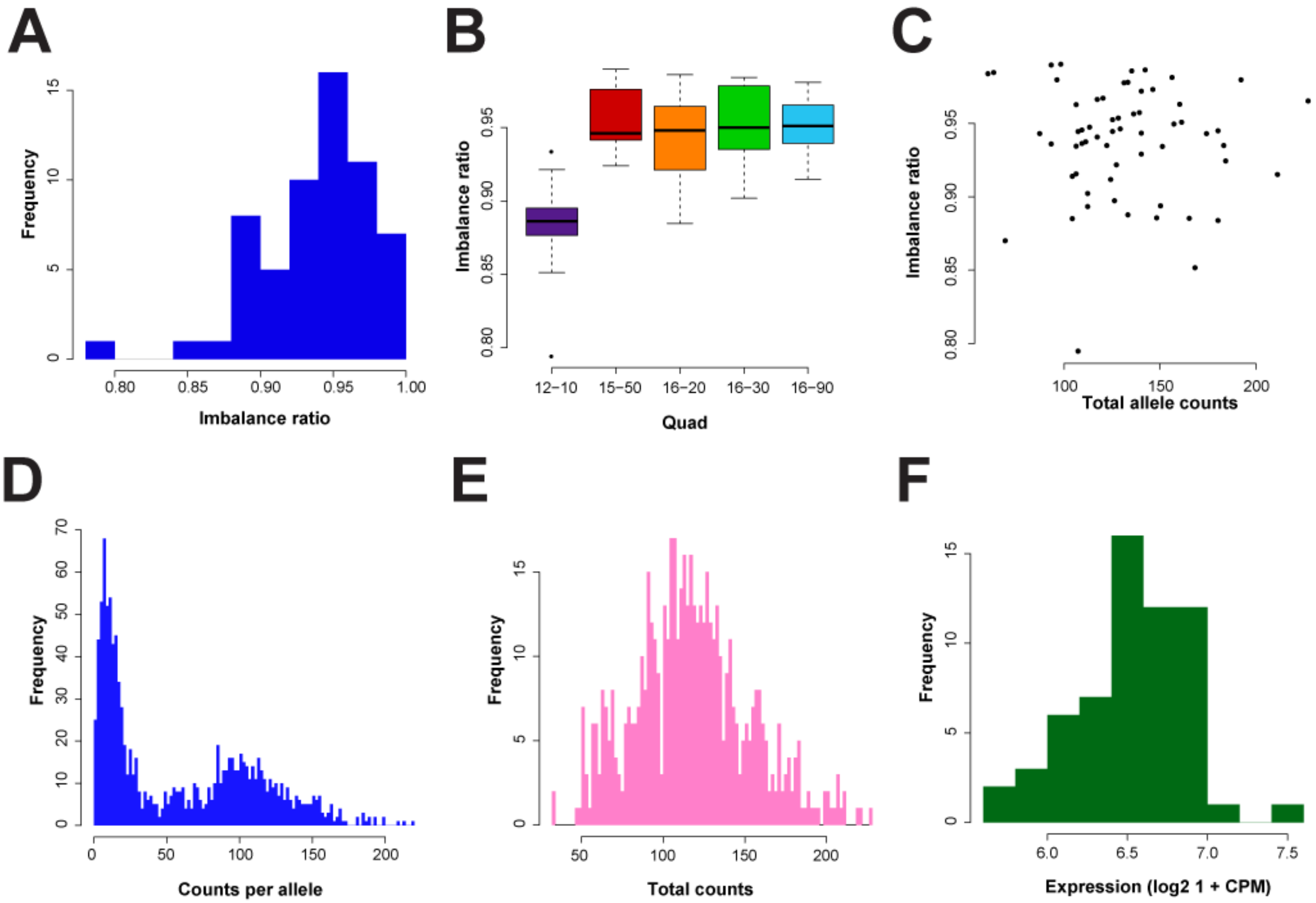

**Fig. S14 Toll-like receptor 1 gene (TLR1) expression:**

*TLR1* is a risk gene for leprosy. Here we show how allele specific expression imbalance does not impact total expression levels, despite one allele being more expressed than the other.

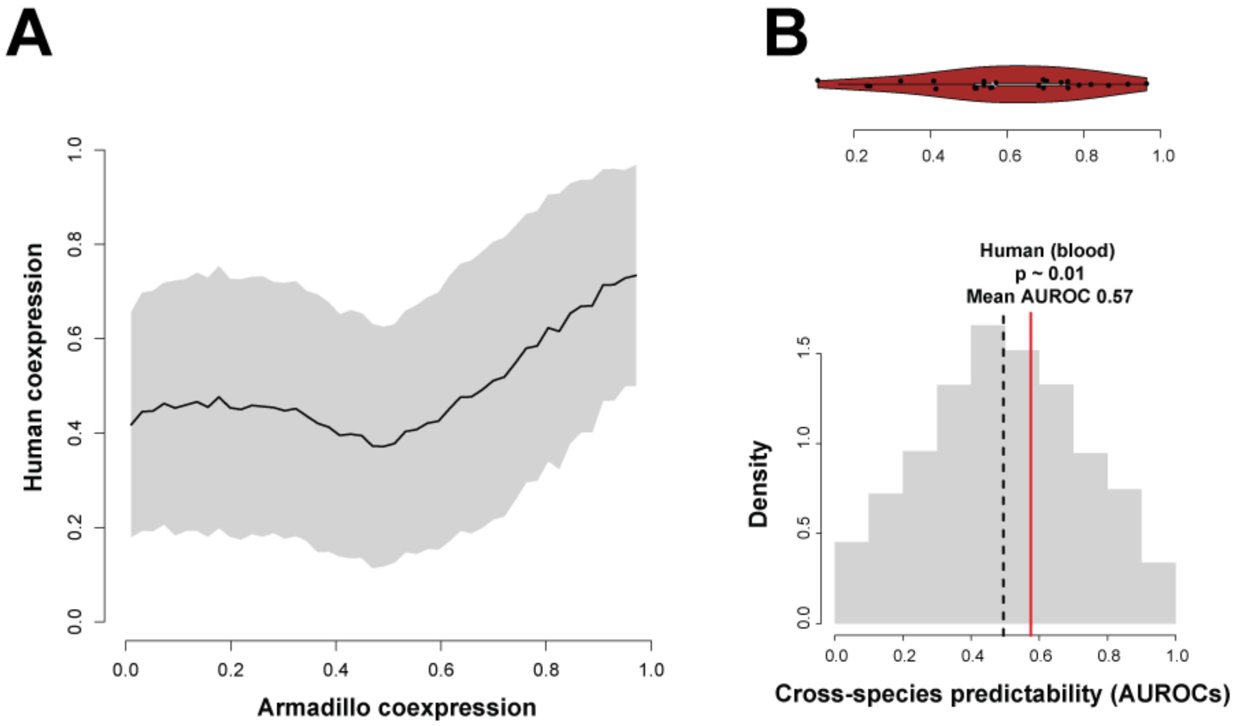

**Fig. S15 Comparing co-expression and cross-species predictability:**

(A) Smoothed co-expression between armadillos and humans across all common gene-pairs (B) The perfect predictor modules perform on average above the null in a human aggregate co-expression network.

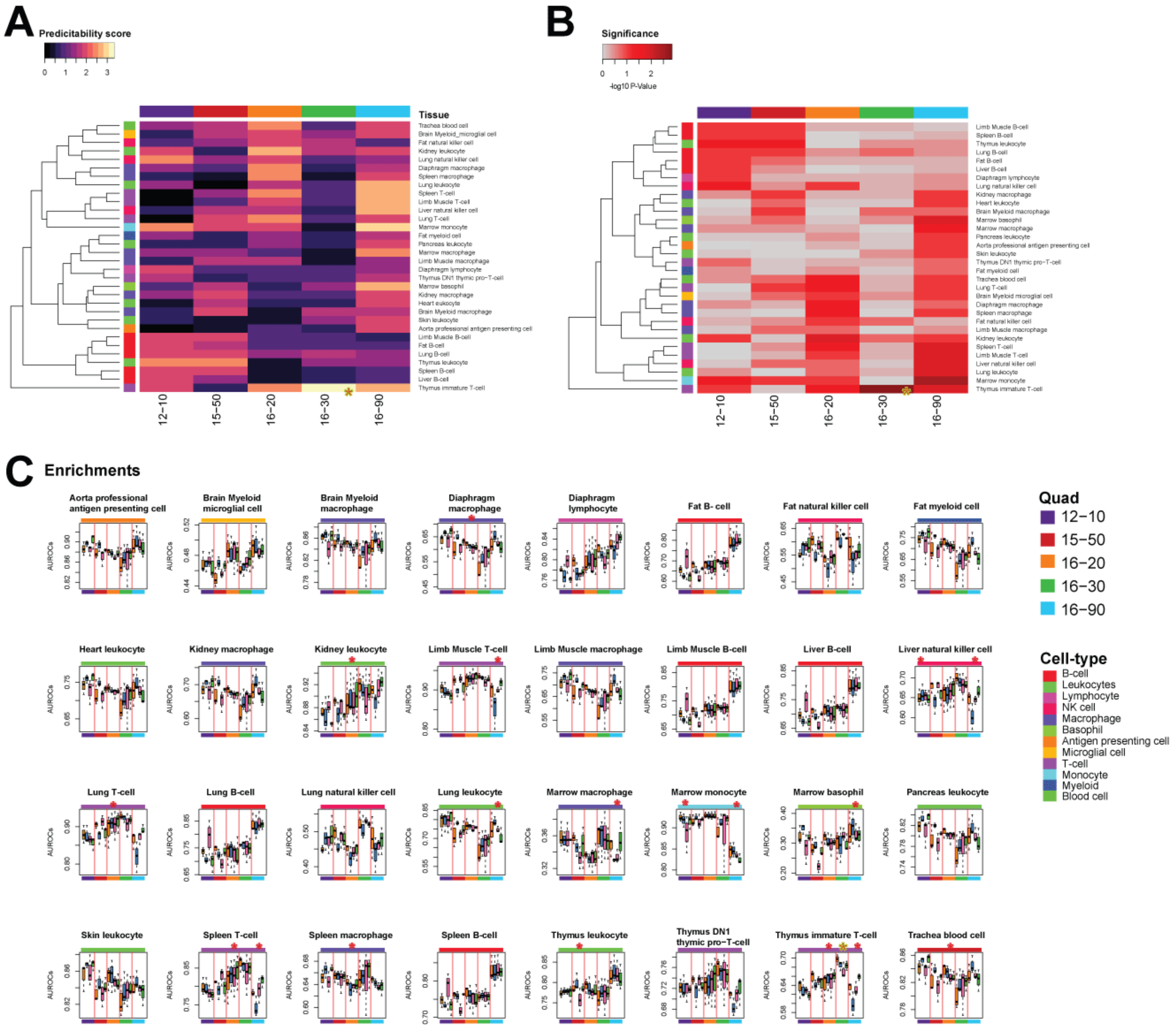

**Fig. S16 Enrichments of cell-type markers from Tabula Muris**

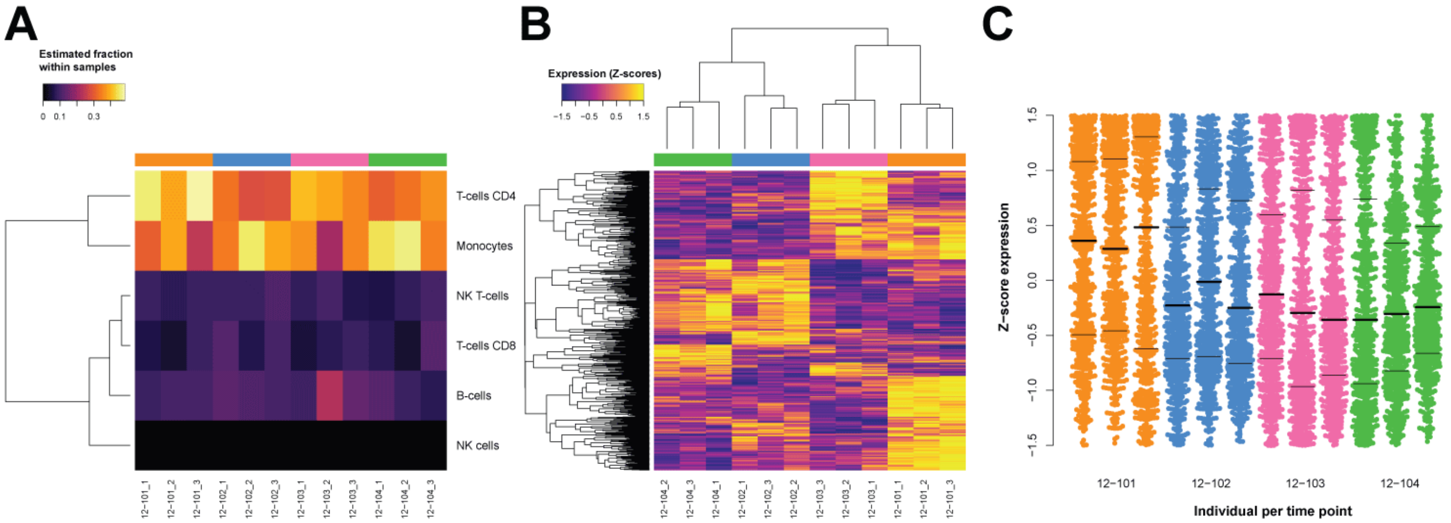

**Fig. S17 Expression properties for quad 12-10:**

(A) Fraction estimates of immune cell types for quad 12-10 using CIBERSORTx. (B-C) Genes most DE via ANOVA for quad 12-10. No genes were significant after correcting for multiple tests. These are based on the unadjusted p-values.

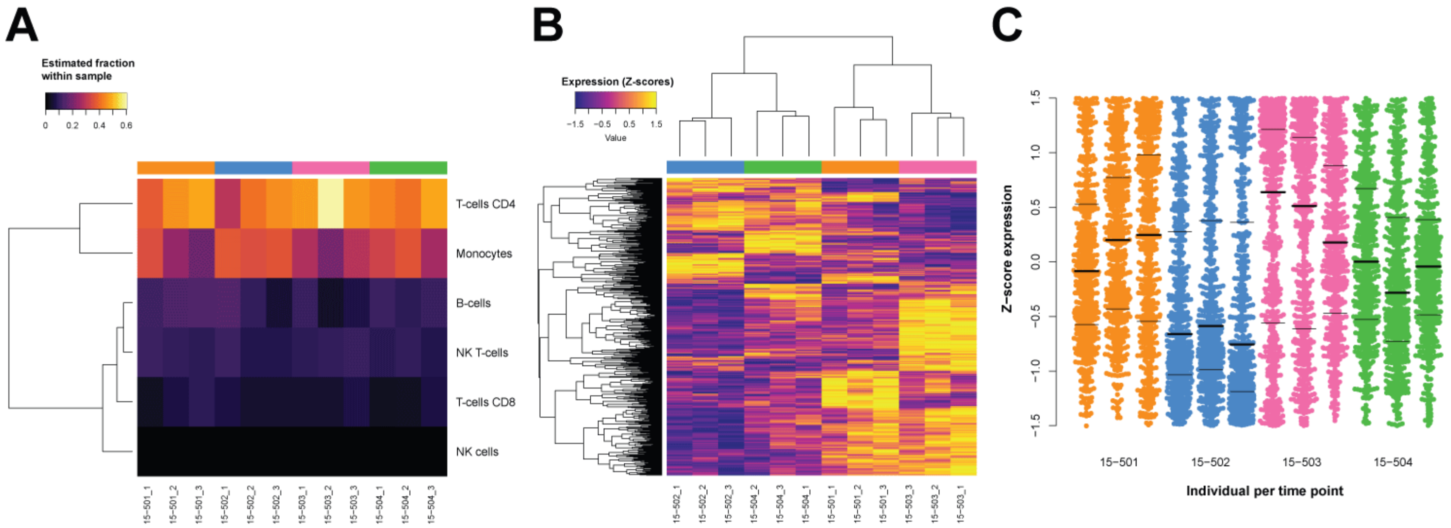

**Fig. S18 Expression properties for quad 15-50:**

(A) Fraction estimates of immune cell types for quad 15-50 using CIBERSORTx. (B-C) Genes most DE via ANOVA for quad 15-50. No genes were significant after correcting for multiple tests. These are based on the unadjusted p-values.

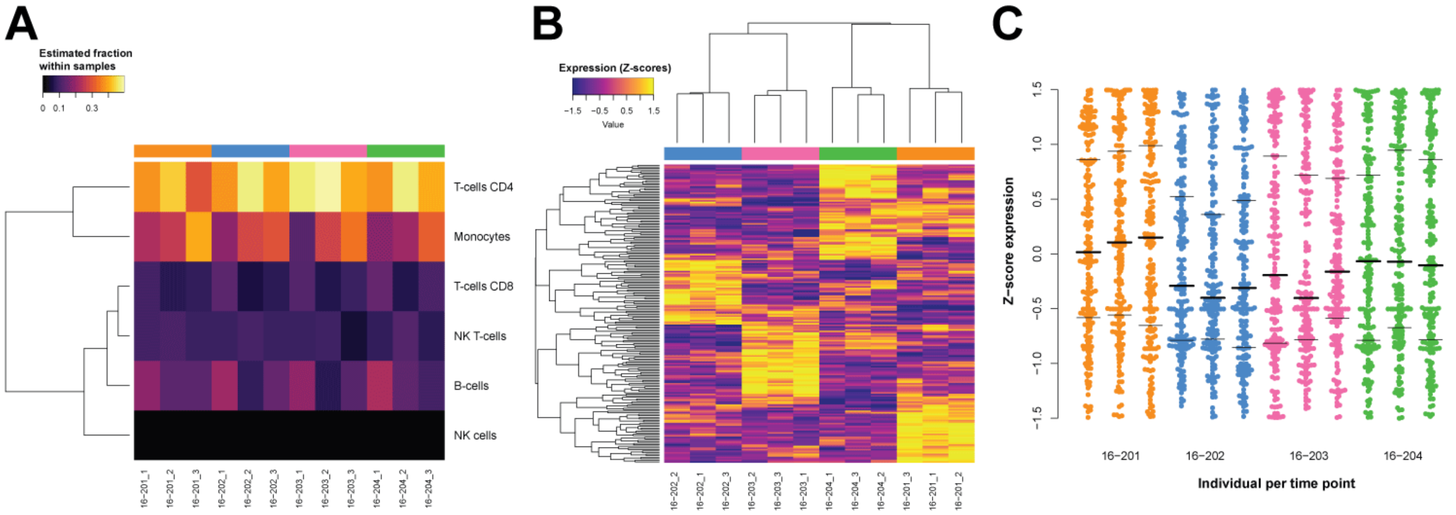

**Fig. S19 Expression properties for quad 16-20:**

(A) Fraction estimates of immune cell types for quad 16-20 using CIBERSORTx. (B-C) Genes most DE via ANOVA for quad 16-20. No genes were significant after correcting for multiple tests. These are based on the unadjusted p-values.

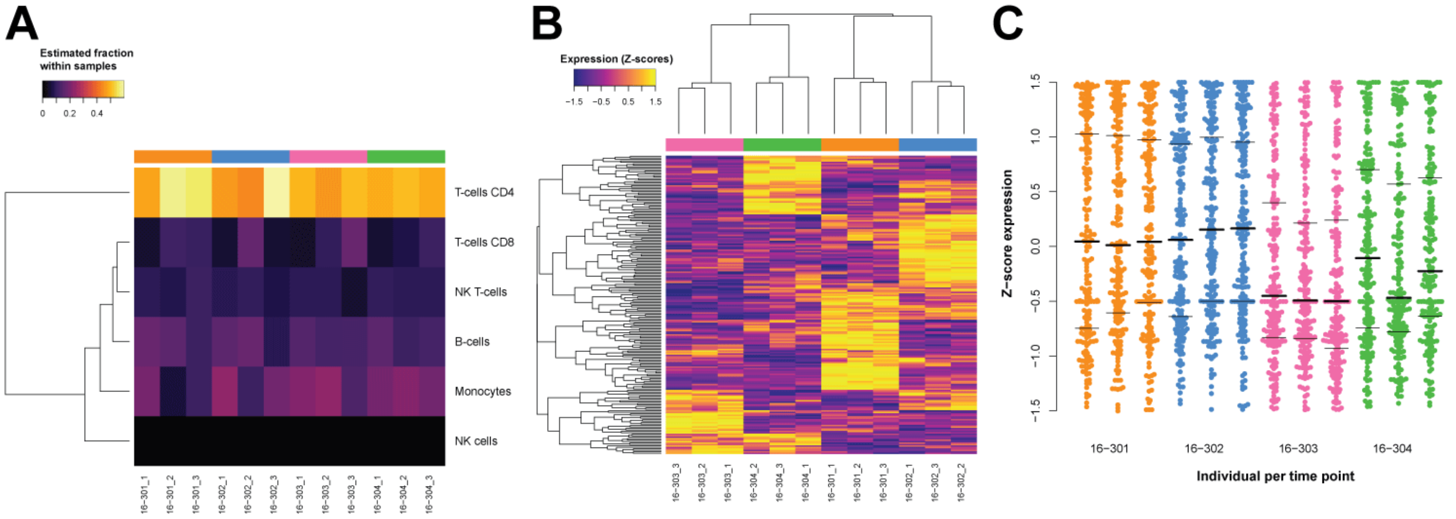

**Fig. S20 Expression properties for quad 16-30:**

(A) Fraction estimates of immune cell types for quad 16-30 using CIBERSORTx. (B-C) Genes most DE via ANOVA for quad 16-30. No genes were significant after correcting for multiple tests. These are based on the unadjusted p-values.

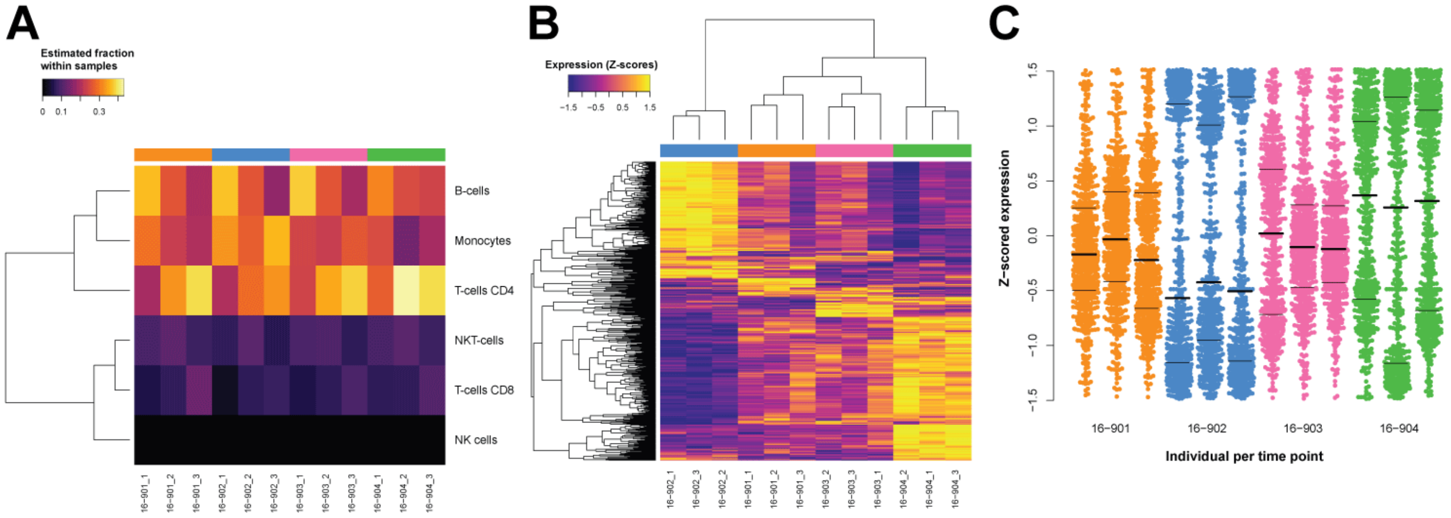

**Fig. S21 Expression properties for quad 16-90:**

(A) Fraction estimates of immune cell types for quad 16-90 using CIBERSORTx. (B-C) Genes most DE via ANOVA for quad 16-90. No genes were significant after correcting for multiple tests. These are based on the unadjusted p-values. However, a majority of genes are under expressed relative to the siblings of individual 16-902.

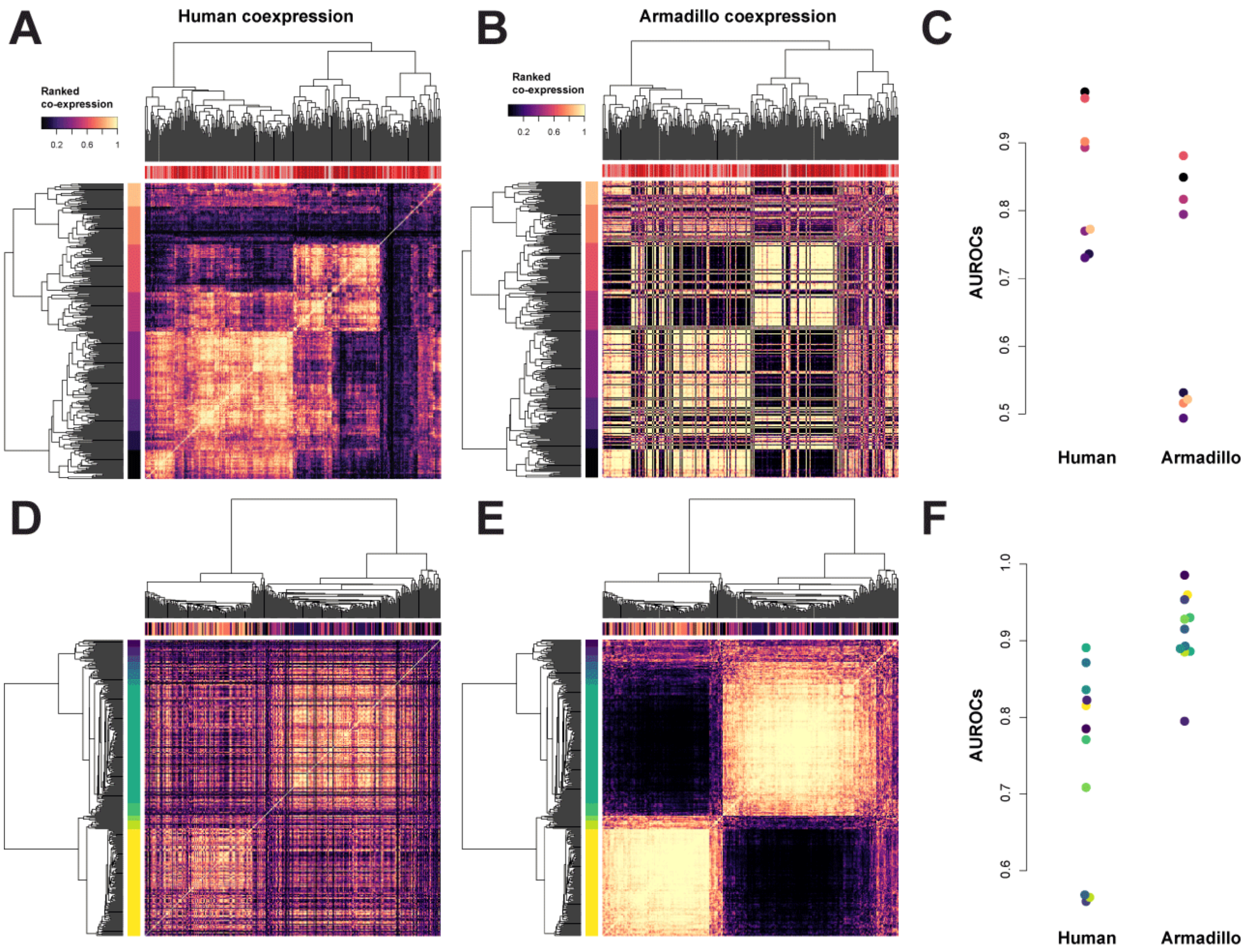

**Fig. S22 Co-expression analysis of quad 16-90:**

(A,D) ANOVA DE genes from quad 16-90 in human and (B,E) armadillo aggregate co-expression networks along with their respective performances based on clustering by human (C) or armadillo (F).

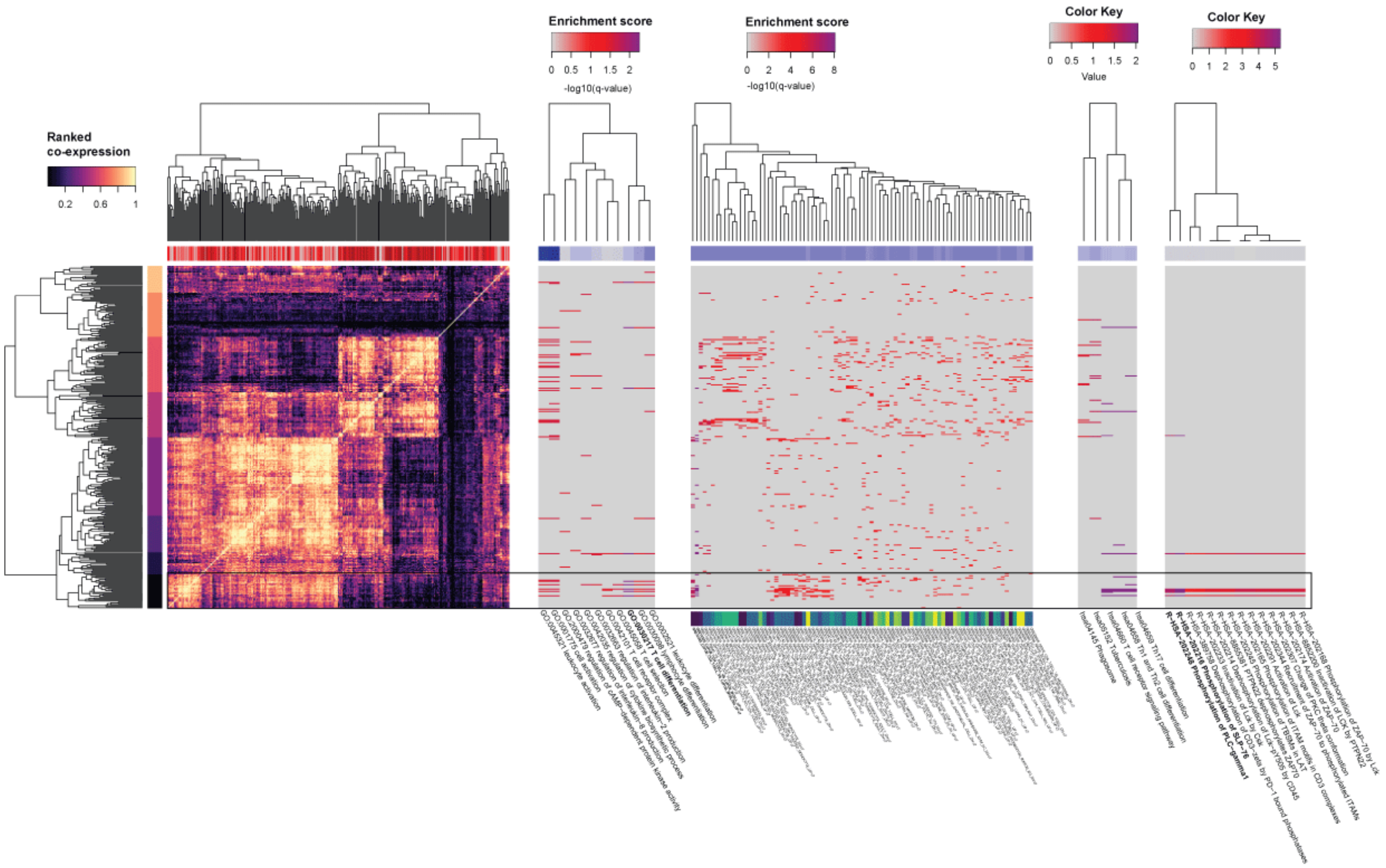

**Fig. S23 Gene set enrichment analysis:**

GO, MSigDB, KEGG, and Reactome pathway enrichments for the ANOVA DE genes. The strongest signal is for T-cell differentiation.

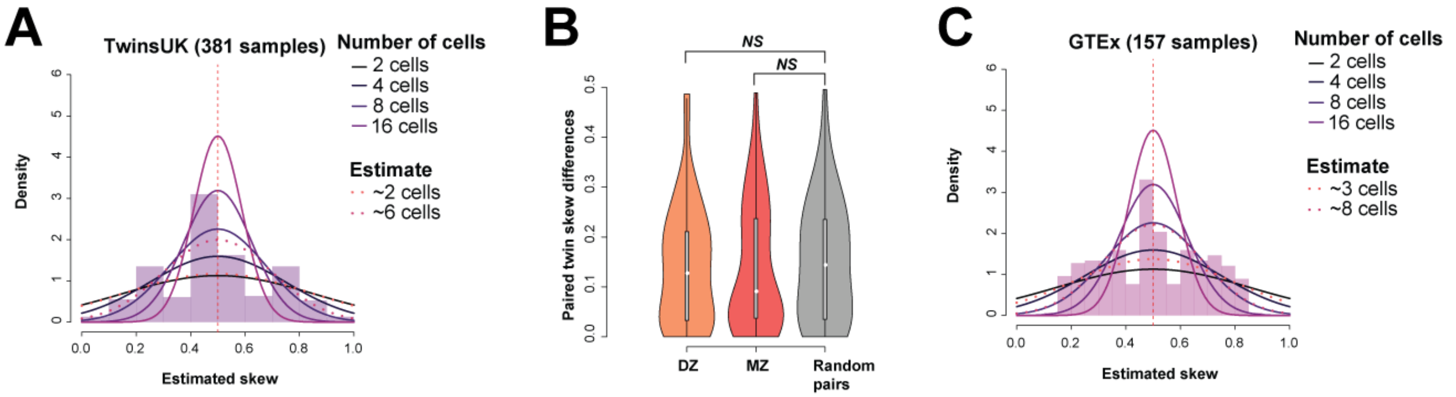

**Fig. S24 Human X-inactivation in blood samples:**

(A) X-inactivation profiles for female blood samples from the TwinsUK data set. (B) There are no significant similarities between skews across twins. (C) GTEx Whole blood samples also shows a similar pattern.

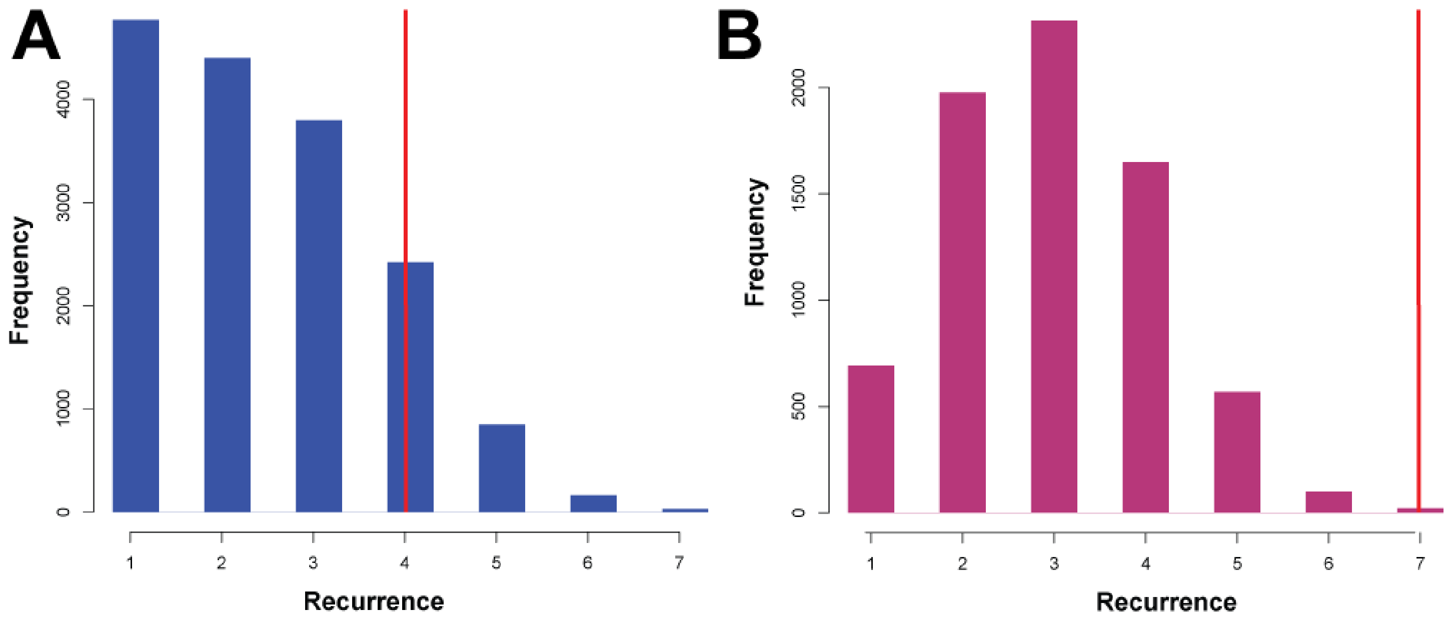

**Fig. S25 eGene recurrence across studies:**

(A) For all genes - over 3000 eGenes are significantly recurrent at an FDR < 0.05 (red lines). (B) For human homologs (7K) – 21 genes are significantly recurrent.

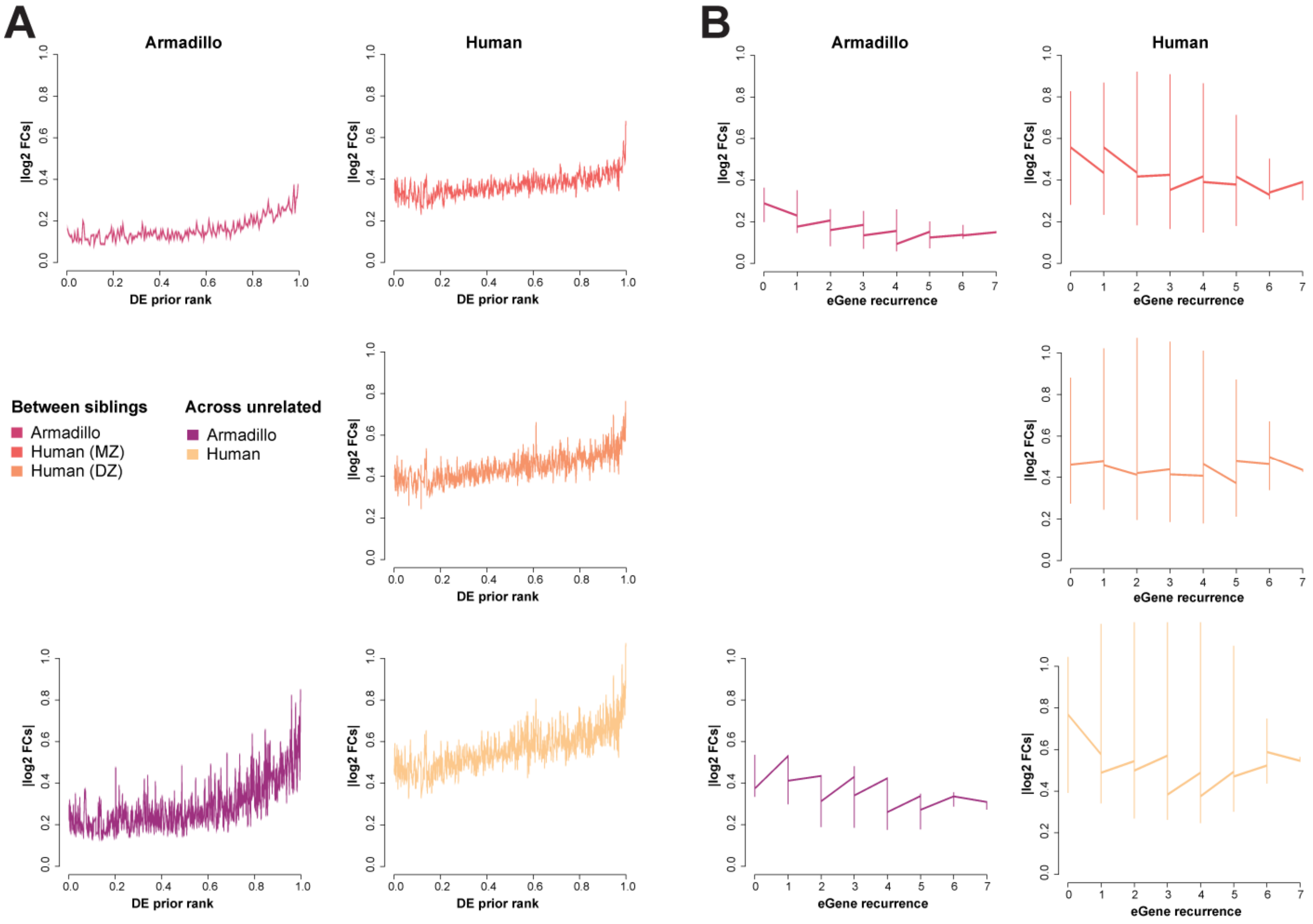

**Fig. S26 Fold change comparisons:**

Comparison of fold changes to DE prior (A) and eGene recurrence (B). The DE prior ranks genes based on their differential expression frequency. eGenes (genes with known cis-eQTLs from blood) show a trend towards lower fold change, suggesting stronger regulatory control (and lower variance).

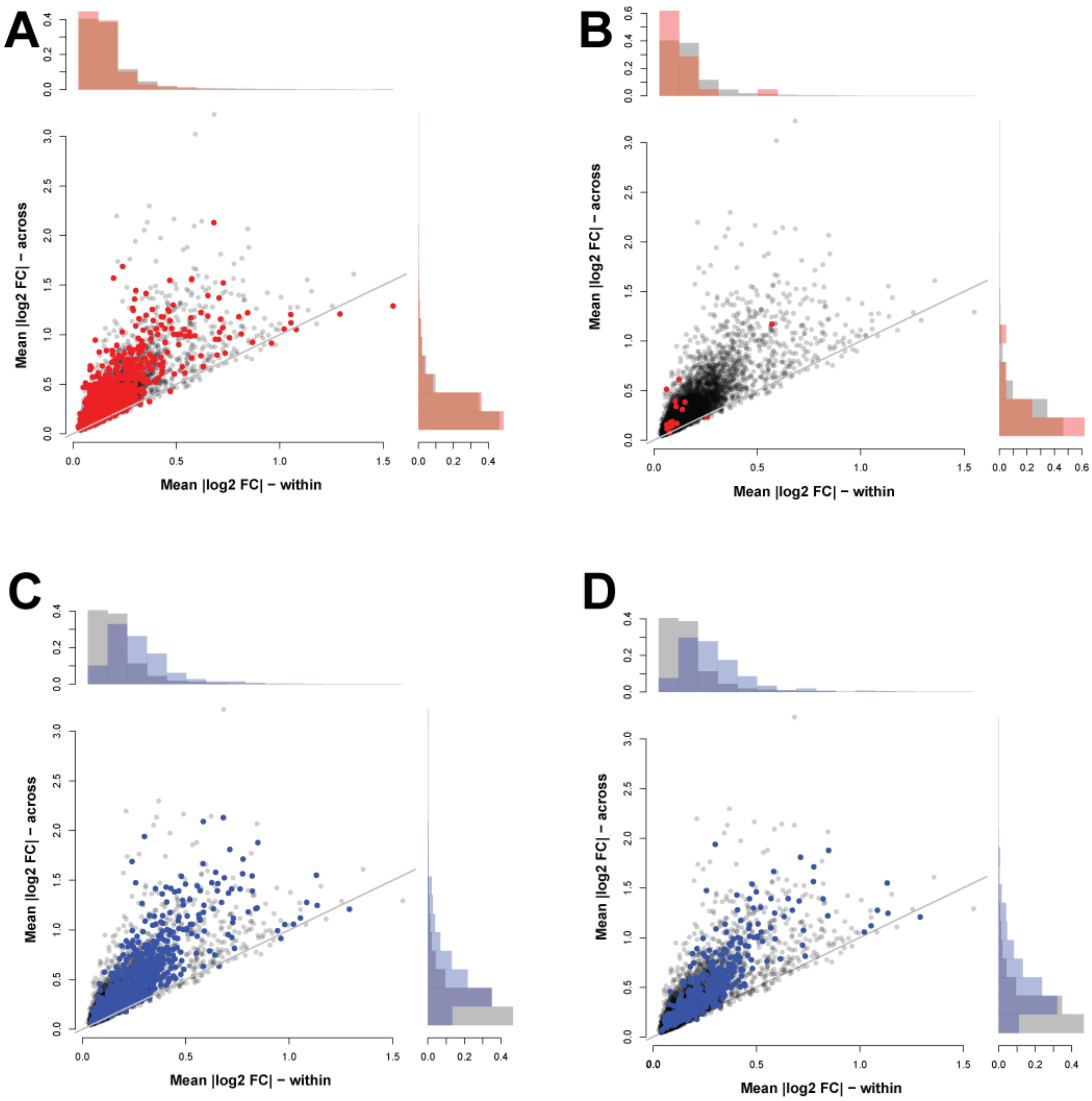

**Fig. S27 Fold change comparisons within and across armadillos:**

(A) eGenes >=4 (B) eGenes >= 7 (C) DE prior rank >= 0.9 (D) DE prior rank >= 0.95.

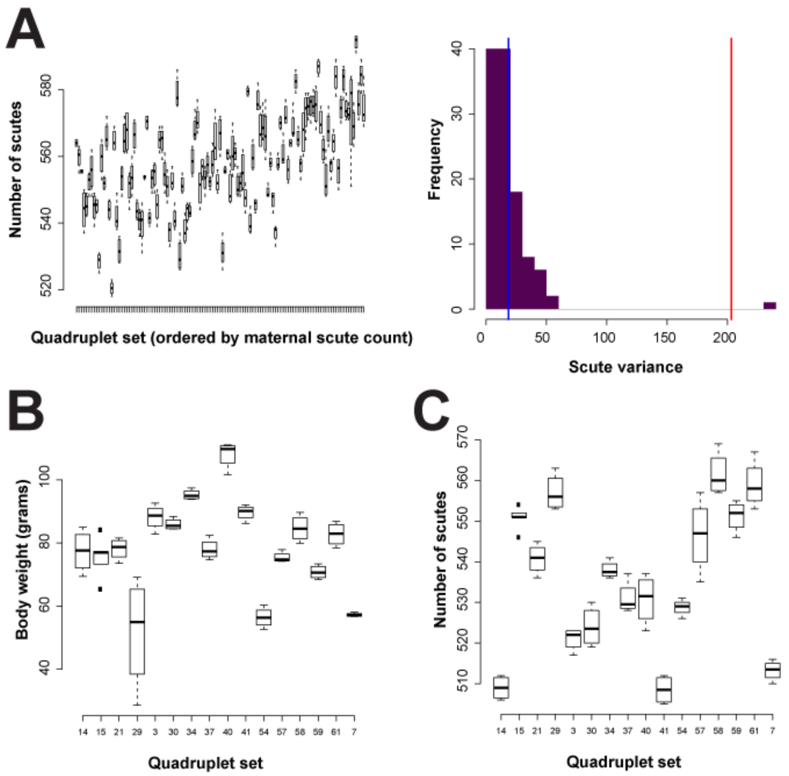

**Fig. S28 Historical phenotypic data showing high similarity within quadruplets:**

(A) Data taken from a collection of 115 (56 male, 59 female) quadruplets from Newman 1913 (*1*) (Table 1). The scutes of the fetuses were counted, as was the mothers. The trend between the number of scutes on the mother and the litter indicates/suggests a genetic component. (D) and (C) are data from 16 litters from Storrs 1967 (*2*) (Table IV).

### Supplementary Tables

**Table S1 Blood collection time points**

| QuadID | Sex | IndividualID | Collected | Received | Age | Collected | Received | Age | Collected | Received | Age |
|---|---|---|---|---|---|---|---|---|---|---|---|
| 12-10 | F | 12-101 | 3/1/2017 | 4/12/2017 | 5 | 2/1/2018 | 3/15/2018 | 6 | 5/10/2018 | 6/26/2018 | 6 |
| 12-10 | F | 12-102 | 3/1/2017 | 4/12/2017 | 5 | 2/1/2018 | 3/15/2018 | 6 | 5/10/2018 | 6/26/2018 | 6 |
| 12-10 | F | 12-103 | 3/1/2017 | 4/12/2017 | 5 | 2/1/2018 | 3/15/2018 | 6 | 5/10/2018 | 6/26/2018 | 6 |
| 12-10 | F | 12-104 | 3/1/2017 | 4/12/2017 | 5 | 2/1/2018 | 3/15/2018 | 6 | 5/10/2018 | 6/26/2018 | 6 |
| 15-50 | M | 15-501 | 3/1/2017 | 4/12/2017 | 2 | 2/1/2018 | 3/15/2018 | 3 | 5/2/2018 | 6/26/2018 | 3 |
| 15-50 | M | 15-502 | 3/1/2017 | 4/12/2017 | 2 | 2/1/2018 | 3/15/2018 | 3 | 5/2/2018 | 6/26/2018 | 3 |
| 15-50 | M | 15-503 | 3/1/2017 | 4/12/2017 | 2 | 2/1/2018 | 3/15/2018 | 3 | 5/2/2018 | 6/26/2018 | 3 |
| 15-50 | M | 15-504 | 3/1/2017 | 4/12/2017 | 2 | 2/1/2018 | 3/15/2018 | 3 | 5/2/2018 | 6/26/2018 | 3 |
| 16-20 | F | 16-201 | 8/1/2017 | 8/18/2017 | 1 | 2/1/2018 | 3/15/2018 | 2 | 7/20/2018 | 9/13/2018 | 2 |
| 16-20 | F | 16-202 | 8/1/2017 | 8/18/2017 | 1 | 2/1/2018 | 3/15/2018 | 2 | 7/20/2018 | 9/13/2018 | 2 |
| 16-20 | F | 16-203 | 8/1/2017 | 8/18/2017 | 1 | 2/1/2018 | 3/15/2018 | 2 | 7/20/2018 | 9/13/2018 | 2 |
| 16-20 | F | 16-204 | 8/1/2017 | 8/18/2017 | 1 | 2/1/2018 | 3/15/2018 | 2 | 7/20/2018 | 9/13/2018 | 2 |
| 16-30 | F | 16-301 | 8/1/2017 | 8/18/2017 | 1 | 2/1/2018 | 3/15/2018 | 2 | 7/20/2018 | 9/13/2018 | 2 |
| 16-30 | F | 16-302 | 8/1/2017 | 8/18/2017 | 1 | 2/1/2018 | 3/15/2018 | 2 | 7/20/2018 | 9/13/2018 | 2 |
| 16-30 | F | 16-303 | 8/1/2017 | 8/18/2017 | 1 | 2/1/2018 | 3/15/2018 | 2 | 7/20/2018 | 9/13/2018 | 2 |
| 16-30 | F | 16-304 | 8/1/2017 | 8/18/2017 | 1 | 2/1/2018 | 3/15/2018 | 2 | 7/20/2018 | 9/13/2018 | 2 |
| 16-90 | M | 16-901 | 8/1/2017 | 8/18/2017 | 1 | 2/1/2018 | 3/15/2018 | 2 | 8/3/2018 | 9/13/2018 | 2 |
| 16-90 | M | 16-902 | 8/1/2017 | 8/18/2017 | 1 | 2/1/2018 | 3/15/2018 | 2 | 8/3/2018 | 9/13/2018 | 2 |
| 16-90 | M | 16-903 | 8/1/2017 | 8/18/2017 | 1 | 2/1/2018 | 3/15/2018 | 2 | 8/3/2018 | 9/13/2018 | 2 |
| 16-90 | M | 16-904 | 8/1/2017 | 8/18/2017 | 1 | 2/1/2018 | 3/15/2018 | 2 | 8/3/2018 | 9/13/2018 | 2 |

**Table S2 RNA-sequencing summary**

| | |
|---|---|
| **Machine** | NextSeq500 |
| **Sequencing** | PE76 |
| | Dual index |
| | KAPA polyA |

| **Flow cell** | HLKF7AFXX | HLJF2AFXX | HHFFKBGX3 | HKNNFBGX5 | HKNNGBGX5 | H7MVWBGX9 | H2LFLBGX9 |
|---|---|---|---|---|---|---|---|
| **Library ID** | 298826 | 298827 | 299555 | 300489 | 300490 | 298975 | 298976 |
| **Run** | Mid Output | Mid Output | High Output | High Output | High Output | High Output | High Output |
| **Date** | 17-05-09 | 17-05-09 | 17-09-15 | 18-03-29 | 18-03-29 | 18-10-19 | 18-10-19 |
| **Clusters (Raw)** | 183029349 | 201345771 | 627521395 | 633345054 | 597025358 | 592280827 | 639878213 |
| **Clusters (PF)** | 165467211 | 179247059 | 569048477 | 565141569 | 545800888 | 518382260 | 540853374 |
| **Yield (MBases)** | 25151 | 27246 | 86495 | 85902 | 82962 | 78794 | 82210 |

**Table S3 Module size and performance of perfect predictors**

| | 12-1 | 15-5 | 16-2 | 16-3 | 16-9 |
|---|---|---|---|---|---|
| **Total number of perfect predictors (genes)** | 65 | 45 | 35 | 29 | 83 |
| **Total number of modules** | 19 | 11 | 20 | 12 | 19 |
| **Modules with at least two genes** | 13 | 8 | 10 | 7 | 9 |
| **Mean AUROCs per** | 0.64 | 0.74 | 0.63 | 0.51 | 0.69 |

**Table S4 Module size and performance of perfect predictors in human co-expression networks**

| | 12-1 | 15-5 | 16-2 | 16-3 | 16-9 |
|---|---|---|---|---|---|
| **Total number of perfect predictors (orthologs)** | 40 | 25 | 20 | 12 | 46 |
| **Total number of modules** | 15 | 7 | 14 | 7 | 12 |
| **Modules with at least two genes** | 9 | 5 | 4 | 2 | 6 |
| **Mean AUROCs – human blood aggregate** | 0.66 | 0.50 | 0.48 | 0.54 | 0.66 |

**Table S5 Human homologs of perfect predictor modules**

| Module | Armadillo | Armadillo, homologs | Human, homologs | Homologs | Gene symbols |
|---|---|---|---|---|---|
| **12-1 2** | **0.50** | 0.76 | **0.82** | 3 | ***SRRM2 PACS2 ACBD4*** |
| **12-1 3** | 0.65 | 0.58 | 0.76 | 5 | *AP5M1 WDR7 EMC2 UBL7 C9orf64* |
| **12-1 4** | 0.75 | 0.40 | 0.79 | 3 | *CDKL5 SPG11 SUFU* |
| **12-1 5** | 0.55 | 0.54 | 0.54 | 7 | *MYBPC2 MYOM1 TIMMDC1 NT5C PLEKHA7 PLEKHG4 NLGN3* |
| **12-1 6** | 0.27 | 0.22 | 0.76 | 2 | *PUM1 SFT2D3* |
| **12-1 7** | 0.81 | 0.78 | 0.57 | 3 | *TM7SF3 FAM45A LRFN4* |
| **12-1 9** | 0.85 | 0.85 | 0.32 | 4 | *CAPZB IFNG HNRNPH2 SMAD1* |
| **12-1 10** | 0.77 | 0.68 | 0.52 | 5 | *PRSS22 ATG4D ST8SIA6 TMEM139 MYZAP* |
| **12-1 11** | **0.94** | 0.96 | **0.86** | 2 | ***NAALADL1 FAM221A*** |
| **15-5 1** | 0.61 | 0.53 | 0.41 | 6 | *ASPM SYT1 SEMA6D ESYT1 ARMC12 ZNF513* |
| **15-5 2** | 0.79 | 0.79 | 0.54 | 10 | *FOXP3 PRPF6 ADAM23 ITGB4 TOMM40L IGFBP6 S100A3 S100A4 TPM2 LRRD1* |
| **15-5 4** | 0.92 | 0.90 | 0.11 | 2 | *KCNMB3 NYNRIN* |
| **15-5 5** | **0.93** | 0.69 | **0.91** | 2 | ***MARC2 SLCO2A1*** |
| **15-5 8** | 0.77 | 0.54 | 0.56 | 3 | *ING3 AGAP2 TRAV41* |
| **16-2 1** | **0.79** | 0.77 | **0.24** | 2 | ***SGSM3 TIMM8B*** |
| **16-2 4** | 0.43 | 0.41 | 0.23 | 2 | *AFAP1 MAK16* |
| **16-2 5** | 0.65 | 0.68 | 0.74 | 3 | *CISH PLCD3 DHRS3* |
| **16-2 6** | 0.52 | 0.54 | 0.69 | 3 | *RSAD2 PGM5 KIAA0895L* |
| **16-3 3** | 0.77 | 0.90 | 0.52 | 5 | *TRIM14 HIP1R PTER ZBTB4 FANCA* |
| **16-3 9** | 0.37 | 0.40 | 0.56 | 2 | *ST6GALNAC2 CERS5* |
| **16-9 1** | 0.85 | 0.83 | 0.68 | 13 | *CACNA1G PTCD2 IL4R GATA3 LCT MPHOSPH10 DDX60 CSGALNACT1 CTLA4 GNL3 RHOH AXIN2 DDX42* |
| **16-9 2** | 0.88 | 0.74 | 0.52 | 3 | *VPS29 UBL3 GATA6* |
| **16-9 3** | 0.39 | 0.48 | 0.96 | 2 | *NALCN SH3PXD2B* |
| **16-9 4** | **0.89** | 0.85 | 0.70 | 7 | ***WWTR1 BIRC3 OSTM1 PPP1R3C XXYLT1 RELL1 FNIP1*** |
| **16-9 6** | 0.88 | 0.82 | 0.69 | 12 | *ASCC2 ETFB SRSF4 IKZF4 PSD4 MDC1 UNC45A USP21 PELO TMC7 SUPT5H NTNG2* |
| **16-9 7** | 0.40 | 0.41 | 0.41 | 3 | *ACADVL MON1A RGMB* |

**Table S6 eQTL and eGene datasets**

| Dataset | Number of eGenes | Reference |
|---|---|---|
| **GEUVADIS – YRI** | 481 | (*3*) |
| **GEUVADIS – EUR** | 3154 | (*3*) |
| **GTEx – blood eQTLs** | 8395 | (*4*) |
| **UCHI_bulk** | 753 | (*5*) |
| **UCHI_mean (single-cell)** | 8428 | (*5*) |
| **condeQTL** | 6664 | (*6*) |
| **bloodeQTL** | 12211 | (*7*) |

**Table S7 Mapping rates to reference genome**

| SampleID | Total reads | Mapped unique | Fraction mapped | SampleID | Total reads | Mapped unique | Fraction mapped |
|---|---|---|---|---|---|---|---|
| 12-101_1 | 41602542 | 36015733 | 86.57% | 15-501_1 | 35282053 | 30339942 | 85.99% |
| 12-101_2 | 54278544 | 47385903 | 87.30% | 15-501_2 | 44446239 | 38451885 | 86.51% |
| 12-101_3 | 40154008 | 34173405 | 85.11% | 15-501_3 | 30792841 | 26110927 | 84.80% |
| 12-102_1 | 38973064 | 32792258 | 84.14% | 15-502_1 | 37266875 | 31888980 | 85.57% |
| 12-102_2 | 46986854 | 40564286 | 86.33% | 15-502_2 | 52735117 | 46173745 | 87.56% |
| 12-102_3 | 34561622 | 29214558 | 84.53% | 15-502_3 | 32965973 | 28177741 | 85.48% |
| 12-103_1 | 37362587 | 32323948 | 86.51% | 15-503_1 | 40360580 | 34979716 | 86.67% |
| 12-103_2 | 48273632 | 41533864 | 86.04% | 15-503_2 | 47881949 | 41387965 | 86.44% |
| 12-103_3 | 37041652 | 30931023 | 83.50% | 15-503_3 | 39966441 | 34243060 | 85.68% |
| 12-104_1 | 38406013 | 33044512 | 86.04% | 15-504_1 | 41064191 | 35589149 | 86.67% |
| 12-104_2 | 53265459 | 45975918 | 86.31% | 15-504_2 | 54408473 | 47865056 | 87.97% |
| 12-104_3 | 39445894 | 33527681 | 85.00% | 15-504_3 | 33643439 | 28342602 | 84.24% |
| 16-201_1 | 45474598 | 39484050 | 86.83% | 16-301_1 | 38922543 | 33780878 | 86.79% |
| 16-201_2 | 45882806 | 40261407 | 87.75% | 16-301_2 | 49669143 | 43797406 | 88.18% |
| 16-201_3 | 34640418 | 29571671 | 85.37% | 16-301_3 | 45974587 | 39159006 | 85.18% |
| 16-202_1 | 39279314 | 34256794 | 87.21% | 16-302_1 | 39645412 | 34175041 | 86.20% |
| 16-202_2 | 42937510 | 38040993 | 88.60% | 16-302_2 | 54415338 | 48152804 | 88.49% |
| 16-202_3 | 37243330 | 31742796 | 85.23% | 16-302_3 | 43267015 | 36715960 | 84.86% |
| 16-203_1 | 47244264 | 41377647 | 87.58% | 16-303_1 | 41475341 | 35384624 | 85.31% |
| 16-203_2 | 48221483 | 42381863 | 87.89% | 16-303_2 | 48007335 | 42685052 | 88.91% |
| 16-203_3 | 42876738 | 36388842 | 84.87% | 16-303_3 | 27402266 | 23012737 | 83.98% |
| 16-204_1 | 43357634 | 37477748 | 86.44% | 16-304_1 | 42662916 | 36925309 | 86.55% |
| 16-204_2 | 54102026 | 47703555 | 88.17% | 16-304_2 | 43762162 | 38625922 | 88.26% |
| 16-204_3 | 41711600 | 35245490 | 84.50% | 16-304_3 | 48557116 | 41159707 | 84.77% |
| 16-901_1 | 52162951 | 43497194 | 83.39% | | | | |
| 16-901_2 | 52082789 | 46009545 | 88.34% | | | | |
| 16-901_3 | 43438263 | 36466131 | 83.95% | | | | |
| 16-902_1 | 41859373 | 35396812 | 84.56% | | | | |
| 16-902_2 | 49328126 | 43342298 | 87.87% | | | | |
| 16-902_3 | 35433898 | 30084997 | 84.90% | | | | |
| 16-903_1 | 44266009 | 37182722 | 84.00% | | | | |
| 16-903_2 | 46358744 | 40791911 | 87.99% | | | | |
| 16-903_3 | 43795015 | 37366506 | 85.32% | | | | |
| 16-904_1 | 38380655 | 32458961 | 84.57% | | | | |
| 16-904_2 | 47565012 | 41494160 | 87.24% | | | | |
| 16-904_3 | 33501327 | 28426115 | 84.85% | | | | |

**Table S8 DNA-sequencing summary from NYGC**

| | |
|---|---|
| **Source Tissue Type** | Blood |
| **Species** | Dasypus novemcinctus |
| **Reference Genome** | Other-non_human_DNA |
| **Library Prep** | TruSeq Nano 450bp |
| **Coverage/Reads** | 30x |

| Quad | DNA collected from | Sex | Volume (µL) | Concentration (ng/µL) | Mass (ng) | GCN |
|---|---|---|---|---|---|---|
| 12-10 | 12D101_12D102_12D103_12D104 | Female | 94 | 17.16 | 1680.09 | 7.7 |
| 15-50 | 15F501_15F502_15F503_15F504 | Male | 94 | 33.71 | 3283.28 | 8.4 |
| 16-20 | 16-201_16-202_16-203_16-204 | Female | 94 | 21.45 | 2099.93 | 7.8 |
| 16-30 | 16-302_16-303_16-304 | Female | 41 | 16.17 | 1579.78 | 8.3 |
| 16-90 | 16-901_16-902_16-903_16-904 | Male | 110 | 27.38 | 2836.50 | 6.0 |

**Table S9 Mapping rates to personal quadruplet genomes.**

| SampleID | Total reads | Mapped unique | Fraction mapped | SampleID | Total reads | Mapped unique | Fraction mapped |
|---|---|---|---|---|---|---|---|
| 12-101_1 | 41602542 | 34731798 | 83.48% | 15-501_1 | 35282053 | 29401615 | 83.33% |
| 12-101_2 | 54278544 | 44553371 | 82.08% | 15-501_2 | 44446239 | 36375942 | 81.84% |
| 12-101_3 | 40154008 | 33130726 | 82.51% | 15-501_3 | 30792841 | 25377955 | 82.42% |
| 12-102_1 | 38973064 | 31773390 | 81.53% | 15-502_1 | 37266875 | 30893161 | 82.90% |
| 12-102_2 | 46986854 | 38314183 | 81.54% | 15-502_2 | 52735117 | 43516277 | 82.52% |
| 12-102_3 | 34561622 | 28499980 | 82.46% | 15-502_3 | 32965973 | 27524332 | 83.49% |
| 12-103_1 | 37362587 | 31397309 | 84.03% | 15-503_1 | 40360580 | 34337702 | 85.08% |
| 12-103_2 | 48273632 | 38664809 | 80.10% | 15-503_2 | 47881949 | 39652206 | 82.81% |
| 12-103_3 | 37041652 | 29667242 | 80.09% | 15-503_3 | 39966441 | 33282624 | 83.28% |
| 12-104_1 | 38406013 | 31937592 | 83.16% | 15-504_1 | 41064191 | 34628513 | 84.33% |
| 12-104_2 | 53265459 | 43318507 | 81.33% | 15-504_2 | 54408473 | 44678125 | 82.12% |
| 12-104_3 | 39445894 | 32561207 | 82.55% | 15-504_3 | 33643439 | 27535651 | 81.85% |
| 16-201_1 | 45474598 | 38365399 | 84.37% | 16-301_1 | 38922543 | 32758685 | 84.16% |
| 16-201_2 | 45882806 | 37553676 | 81.85% | 16-301_2 | 49669143 | 41658436 | 83.87% |
| 16-201_3 | 34640418 | 28386059 | 81.94% | 16-301_3 | 45974587 | 38107943 | 82.89% |
| 16-202_1 | 39279314 | 33375526 | 84.97% | 16-302_1 | 39645412 | 33504750 | 84.51% |
| 16-202_2 | 42937510 | 35557003 | 82.81% | 16-302_2 | 54415338 | 46100056 | 84.72% |
| 16-202_3 | 37243330 | 30644433 | 82.28% | 16-302_3 | 43267015 | 35661945 | 82.42% |
| 16-203_1 | 47244264 | 40591140 | 85.92% | 16-303_1 | 41475341 | 34423491 | 83.00% |
| 16-203_2 | 48221483 | 40302268 | 83.58% | 16-303_2 | 48007335 | 40759874 | 84.90% |
| 16-203_3 | 42876738 | 35112101 | 81.89% | 16-303_3 | 27402266 | 22311426 | 81.42% |
| 16-204_1 | 43357634 | 36513505 | 84.21% | 16-304_1 | 42662916 | 36066795 | 84.54% |
| 16-204_2 | 54102026 | 45112901 | 83.38% | 16-304_2 | 43762162 | 36821177 | 84.14% |
| 16-204_3 | 41711600 | 34172098 | 81.92% | 16-304_3 | 48557116 | 40042965 | 82.47% |
| 16-901_1 | 52162951 | 42319360 | 81.13% | | | | |
| 16-901_2 | 52082789 | 43831059 | 84.16% | | | | |
| 16-901_3 | 43438263 | 35135909 | 80.89% | | | | |
| 16-902_1 | 41859373 | 34667948 | 82.82% | | | | |
| 16-902_2 | 49328126 | 40601505 | 82.31% | | | | |
| 16-902_3 | 35433898 | 28899493 | 81.56% | | | | |
| 16-903_1 | 44266009 | 36313376 | 82.03% | | | | |
| 16-903_2 | 46358744 | 38302650 | 82.62% | | | | |
| 16-903_3 | 43795015 | 36015299 | 82.24% | | | | |
| 16-904_1 | 38380655 | 31433631 | 81.90% | | | | |
| 16-904_2 | 47565012 | 39127091 | 82.26% | | | | |
| 16-904_3 | 33501327 | 27349728 | 81.64% | | | | |

**Table S10 Reciprocal top BLAST hits for *XIST* in humans**

| Armadillo ID | Top human hit | Gene | Sequence length | Alignment |
|---|---|---|---|---|
| ENSDNOG00000000718 | CHR_HSCHR10_1_CTG4:77821512-77822104 [Sequence] | DLG5 | 593 [Sequence] | 86.34 [Alignment] |
| ENSDNOG00000032486 | 14:49586580-49586872 [Sequence] | RPS29, AL139099.5, RN7SL1 | 293 [Sequence] | 90.10 [Alignment] |
| ENSDNOG00000033080 | X:73831145-73831260 [Sequence] | XIST, Xist_exon4 | 116 [Sequence] | 95.69 [Alignment] |
| ENSDNOG00000033288 | 10:42566677-42567476 [Sequence] | EIF3LP2 | 801 [Sequence] | 89.64 [Alignment] |
| ENSDNOG00000034036 | X:120691217-120691290 [Sequence] | | 74 [Sequence] | 93.24 [Alignment] |
| | 14:74126451-74126520 [Sequence] | LIN52 | 70 [Sequence] | 94.29 [Alignment] |
| ENSDNOG00000034452 | 14:49586580-49586879 [Sequence] | RPS29, AL139099.5, RN7SL1 | 300 [Sequence] | 98.33 [Alignment] |
| ENSDNOG00000035926 | X:55741080-55741563 [Sequence] | RRAGB | 484 [Sequence] | 86.36 [Alignment] |
| ENSDNOG00000036040 | 11:5453493-5454420 [Sequence] | HBG2, HBE1, AC087380.1, OR51I2 | 928 [Sequence] | 86.96 [Alignment] |
| ENSDNOG00000037020 | 11:5544577-5545088 [Sequence] | HBG2, OR52H1 | 512 [Sequence] | 87.70 [Alignment] |
| ENSDNOG00000039691 | 6:134133592-134133852 [Sequence] | RN7SL408P | 261 [Sequence] | 88.89 [Alignment] |
| ENSDNOG00000040606 | 3:11103033-11103275 [Sequence] | | 243 [Sequence] | 72.84 [Alignment] |
| ENSDNOG00000042044 | 14:49862649-49862841 [Sequence] | AL627171.2, RN7SL2 | 193 [Sequence] | 93.78 [Alignment] |
| ENSDNOG00000042440 | 14:49586588-49586780 [Sequence] | RPS29, AL139099.5, RN7SL1 | 193 [Sequence] | 93.78 [Alignment] |

| Armadillo ID | Top human hit | Gene | Sequence length | Alignment |
|---|---|---|---|---|
| | X:140085873-140086037 [Sequence ] | RN7SL727P | 165 [Sequence] | 89.09 [Alignment] |
| ENSDNOG00000042960 | 3:15738515-15738715 [Sequence] | ANKRD28, RN7SL4P | 201 [Sequence] | 94.53 [Alignment] |
| ENSDNOG00000045206 | 16:71137144-71137399 [Sequence] | HYDIN | 256 [Sequence] | 88.67 [Alignment] |
| ENSDNOG00000047775 | X:73821657-73821724 [Sequence] | TSIX, XIST, XIST_intron | 68 [Sequence] | 92.65 [Alignment] |

**Table S11 Reciprocal top BLAST hits for *XIST* in mouse**

| Armadillo ID | Top mouse hit | Gene | Sequence length | Alignment |
|---|---|---|---|---|
| ENSDNOG00000000718 | 14:24158165-24158756 [Sequence] | Dlg5 | 592 [Sequence] | 79.90 [Alignment] |
| ENSDNOG00000032486 | 12:69159295-69159587 [Sequence] | Rn7s1, AC099934.2, AC099934.1 | 293 [Sequence] | 89.42 [Alignment] |
| ENSDNOG00000033080 | X:103469677-103469758 [Sequence] | Gm26992, Tsix, Xist, Gm27927 | 82 [Sequence] | 93.90 [Alignment] |
| ENSDNOG00000033288 | 15:79089463-79089960 [Sequence] | Eif3l | 498 [Sequence] | 88.96 [Alignment] |
| ENSDNOG00000034036 | 18:80197662-80197795 [Sequence] | Rbfa | 134 [Sequence] | 85.82 [Alignment] |
| | X:155157596-155157655 [Sequence] | | 60 [Sequence] | 91.67 [Alignment] |
| ENSDNOG00000034452 | 12:69159295-69159594 [Sequence] | Rn7s1, AC099934.2, AC099934.1 | 300 [Sequence] | 98.33 [Alignment] |
| ENSDNOG00000035926 | 14:24164394-24164579 [Sequence] | Dlg5 | 186 [Sequence] | 89.25 [Alignment] |
| ENSDNOG00000036040 | 7:104049443-104050316 [Sequence] | Olfr643, Olfr642 | 875 [Sequence] | 85.60 [Alignment] |
| ENSDNOG00000037020 | 7:104049443-104050316 [Sequence] | Olfr643, Olfr642 | 875 [Sequence] | 85.60 [Alignment] |
| ENSDNOG00000039691 | 12:69159400-69159583 [Sequence] | Rn7s1, AC099934.2, AC099934.1 | 184 [Sequence] | 96.20 [Alignment] |
| ENSDNOG00000040606 | 12:63684161-63684362 [Sequence] | | 202 [Sequence] | 72.77 [Alignment] |
| ENSDNOG00000042044 | 6:69516344-69516539 [Sequence] | Rn7s6, AC156953.1 | 196 [Sequence] | 92.86 [Alignment] |
| ENSDNOG00000042440 | 12:69159314-69159495 [Sequence] | Rn7s1, AC099934.2 | 182 [Sequence] | 94.51 [Alignment] |
| | X:87899315-87899489 [Sequence] | Il1rapl1, Gm24812 | 175 [Sequence] | 84.00 [Alignment] |
| ENSDNOG00000042960 | 6:69516339-69516539 [Sequence] | Rn7s6, AC156953.1 | 201 [Sequence] | 94.03 [Alignment] |
| ENSDNOG00000045206 | 6:131253488-131253758 [Sequence] | Gm5582 | 276 [Sequence] | 80.43 [Alignment] |

| Armadillo ID | Top mouse hit | Gene | Sequence length | Alignment |
|---|---|---|---|---|
| ENSDNOG00000047775 | X:103461220-103461264 [Sequence] | Gm26992, Tsix, Xist, Gm27733 | 45 [Sequence] | 88.89 [Alignment] |
